# p140Cap enhances breast cancer chemosensitivity by limiting an ABCC1-enriched stem-like compartment via β-Catenin inhibition

**DOI:** 10.64898/2026.08.14.744806

**Authors:** Andrea Scavuzzo, Matteo Poncina, Alessia Lamolinara, Alessandro Sarcinella, Mahdis Jahanbin, Maria Grazia Filippone, Laura Bottoni, Francesco Antonio Tucci, Yaron Vinik, Sima Lev, Manuela Iezzi, Ugo Ala, Daniela Taverna, Francesca Orso, Barbara Belletti, Emilia Turco, Salvatore Pece, Daniela Tosoni, Paola Defilippi, Vincenzo Salemme

**Affiliations:** Department of Molecular Biotechnology and Health Sciences, University of Torino, Via Nizza 52, 10126 Torino, Italy; Molecular Biotechnology Center (MBC) “Guido Tarone”, Via Nizza, 52, 10126 Turin, Italy; Agora Cancer Research Center, Lausanne, Switzerland; Immuno-Oncology Laboratory, Center for Advanced Studies and Technology (CAST), Department of Neuroscience, Imaging and Clinical Sciences, G. d’Annunzio University of Chieti-Pescara, Chieti-Pescara, Italy; Department of Medical Sciences, University of Turin, Corso Dogliotti 14, Turin, Italy; Department of Veterinary Sciences, University of Torino, Torino, Italy; European Institute of Oncology IRCCS, 20141 Milan, Italy; Department of Oncology and Hemato-Oncology, Università degli Studi di Milano, 20142 Milano, Italy; Molecular Cell Biology Department, Weizmann Institute of Science, Rehovot, 76100, Israel; Department of Translational Medicine (DIMET), University of Eastern Piedmont, Via Solaroli 17, Novara, 28100, Italy; Molecular Oncology Unit, Centro di Riferimento Oncologico di Aviano (CRO) IRCCS, National Cancer Institute, Aviano (PN), Italy

## Abstract

Chemotherapy response in breast cancer is highly heterogeneous and influenced by tumor-intrinsic drivers of drug sensitivity, including cancer stem cell abundance. We previously reported that the scaffold protein p140Cap limits breast cancer stem cell traits and delays tumor progression. Here, we investigated the role of p140Cap in shaping sensitivity to chemotherapy in HER2-positive and triple-negative breast cancer. In preclinical and patient-derived models, p140Cap enhances chemotherapy response by increasing intracellular doxorubicin retention, DNA damage and subsequent apoptosis. Mechanistically, p140Cap constrained a doxorubicin-negative side population enriched for stem-like properties and elevated ABCC1 expression via inhibition of β-Catenin signaling. Constitutively active β-Catenin expression reversed this phenotype, whereas pharmacological inhibition of the Wnt/β-Catenin pathway with IWR-1 or LGK-974 sensitized p140Cap-deficient tumors to chemotherapy. Clinically, analyses of breast cancer cohorts and patient-derived xenograft models identify p140Cap as predictive biomarker of chemotherapy response, proposing p140Cap-guided patient stratification, dose optimization and rational combination therapies.

## Main

Breast cancer (BC) is the most diagnosed malignancy in women and a leading cause of cancer mortality^1^. Despite advances in subtype-specific targeted therapies, cytotoxic chemotherapy remains the backbone of treatment for high-risk BC^2,3^. In HER2-positive disease, it is routinely combined with targeted agents^4,5^, whereas in triple-negative breast cancer (TNBC), it remains the main therapeutic option and is increasingly being combined with immunotherapy in advanced settings^6–9^. Chemotherapy is widely used but associated with significant toxicities and heterogeneous responses across BC^10^, underscoring the need for molecular strategies that enhance sensitivity and enable precision medicine approaches to optimize treatment intensity and reduce side effects. A major barrier to effective chemotherapy is the presence of breast cancer stem cells (BCSCs), a tumor cell compartment characterized by adaptive traits that shape therapeutic response. These include enhanced drug efflux via ATP-binding cassette (ABC) transporters, such as ABCC1, ABCB1, and ABCG2, which reduce intracellular drug accumulation and thereby attenuate treatment efficacy^11–13^.

BCSCs are closely associated with Wnt/β-Catenin signaling, which supports self-renewal, epithelial-to-mesenchymal transition and metastatic potential across multiple tumor types^14–16^. Therapeutic strategies targeting this pathway have therefore emerged as a promising approach to enhance chemotherapy sensitivity, prevent relapse and improve treatment stratification while reducing toxicity^17–20^. The discovery of novel biomarkers is therefore fundamental to achieve a deeper understanding of the molecular mechanisms that underpin therapy response in BC^21^. Such biomarkers hold the potential to facilitate the development of more personalized treatment strategies, ultimately leading to improved diagnostic and prognostic accuracy, enhanced therapeutic efficacy and a reduction in the adverse effects and economic burden associated with ineffective interventions^22–24^.

In pursuit of dynamic predictors of treatment outcome, we previously identified p140Cap, a scaffold protein encoded by *SRCIN1* gene, as a suppressor of BC progression^25–28^. p140Cap expression correlates with favourable clinicopathological features and reduced metastatic potential and limits the BCSC-like compartment through stabilization of the β-Catenin destruction complex^29^. Defining whether p140Cap shapes chemotherapy responsiveness and may serve as a predictive biomarker of therapeutic response therefore represents the aim of this study.

Here, using HER2 and TNBC cell and preclinical models, we show that p140Cap presence enhances chemotherapy responsiveness. This heightened sensitivity results from p140Cap-mediated shrinkage of the BCSC compartment, which is associated with reduced *ABCC1* expression and consequent increases in intracellular drug retention and apoptosis. Mechanistically, the reduction of BCSC pool is mediated by p140Cap-dependent inhibition of β-Catenin signaling^29^. Consistently, pharmacological inhibition of the β-Catenin pathway phenocopied p140Cap function and restored doxorubicin sensitivity in p140Cap-deficient tumours, supporting Wnt/β-Catenin pathway targeting as a potential combination strategy to improve chemotherapy response in patients with absent p140Cap expression. Finally, analyses of patient cohorts and patient-derived xenografts (PDXs) identify p140Cap as a candidate predictive biomarker of chemotherapy response, with potential translational relevance for patient stratification and therapy optimization.

## Results

### p140Cap enhances chemosensitivity in breast cancer

CSCs limit chemotherapeutic efficacy and drive tumor recurrence and metastatization in BC and most solid tumors^11^. Building on our previous finding that p140Cap reduces the BCSC compartment^29^, we first investigated whether p140Cap expression correlates with treatment outcome in BC patients. Analysis of an independent BC cohort from ROCplot.com^30^ revealed that *SRCIN1* expression is significantly enriched in tumors from patients who responded to chemotherapy treatment compared with non-responders (Fig. 1a). In line with this observation, Receiver Operating Characteristic (ROC) analysis yielded an area under the curve (AUC) of 0.604, supporting an association between *SRCIN1* expression levels and chemotherapy response (Extended Data Fig. 1a). Consistently, patients with high p140Cap expression displayed improved overall survival after neoadjuvant chemotherapy in an integrated METABRIC, TCGA and IMPACT dataset^31^ (Fig. 1b). Although these associations are not indicative of causality, they collectively support a potential link between p140Cap expression and increased chemotherapy sensitivity. To further investigate these observations, we focused on HER2-positive and TNBC subtypes in which chemotherapy constitutes a major therapeutic strategy (Extended Data Fig. 1b)^32^, using the following BC cell models: (i) 4T1, a highly aggressive stage IV murine TNBC; (ii) TuBo, an established HER2⁺ BC cell line derived from BALB/c-MMTV-NeuT mice^33^; (iii) MDA-MB-231, a poorly differentiated model of human TNBC, in which we stably expressed p140Cap or control vector (Mock). These cells were treated for 48 hours with increasing concentrations of Doxorubicin (Doxo)^34^, Paclitaxel^35^ and Vinorelbine^36^, and cell viability was monitored by IncuCyte live-cell imaging. Across all three BC cell lines, p140Cap cells showed reduced cell confluence and approximately 50% decrease in half-maximal inhibitory concentration (IC_50_) values compared with controls (Fig. 1c and Extended Data Fig. 1c). For clinical relevance and experimental tractability, we focused subsequent analyses on doxorubicin, a backbone of BC therapy whose intrinsic red fluorescence further enables direct and quantitative assessment of drug uptake and response^34,37^. To determine whether reduced confluence reflects doxorubicin-induced cell death, apoptosis was quantified by flow cytometric analysis of Annexin V staining, revealing increased apoptotic rate in p140Cap cells after 48 hours of exposure (Fig. 1d). Notably, doubling the doxorubicin dose in control 4T1 cells reduced viability to levels reached by p140Cap-expressing cells treated at half dose (Fig. 1e). Collectively, these results indicate that p140Cap enhances the sensitivity of murine and human BC tumor cells to chemotherapy, suggesting the possibility of effective treatment at lower drug doses compared with p140Cap-deficient breast tumors.

**Fig. 1:**
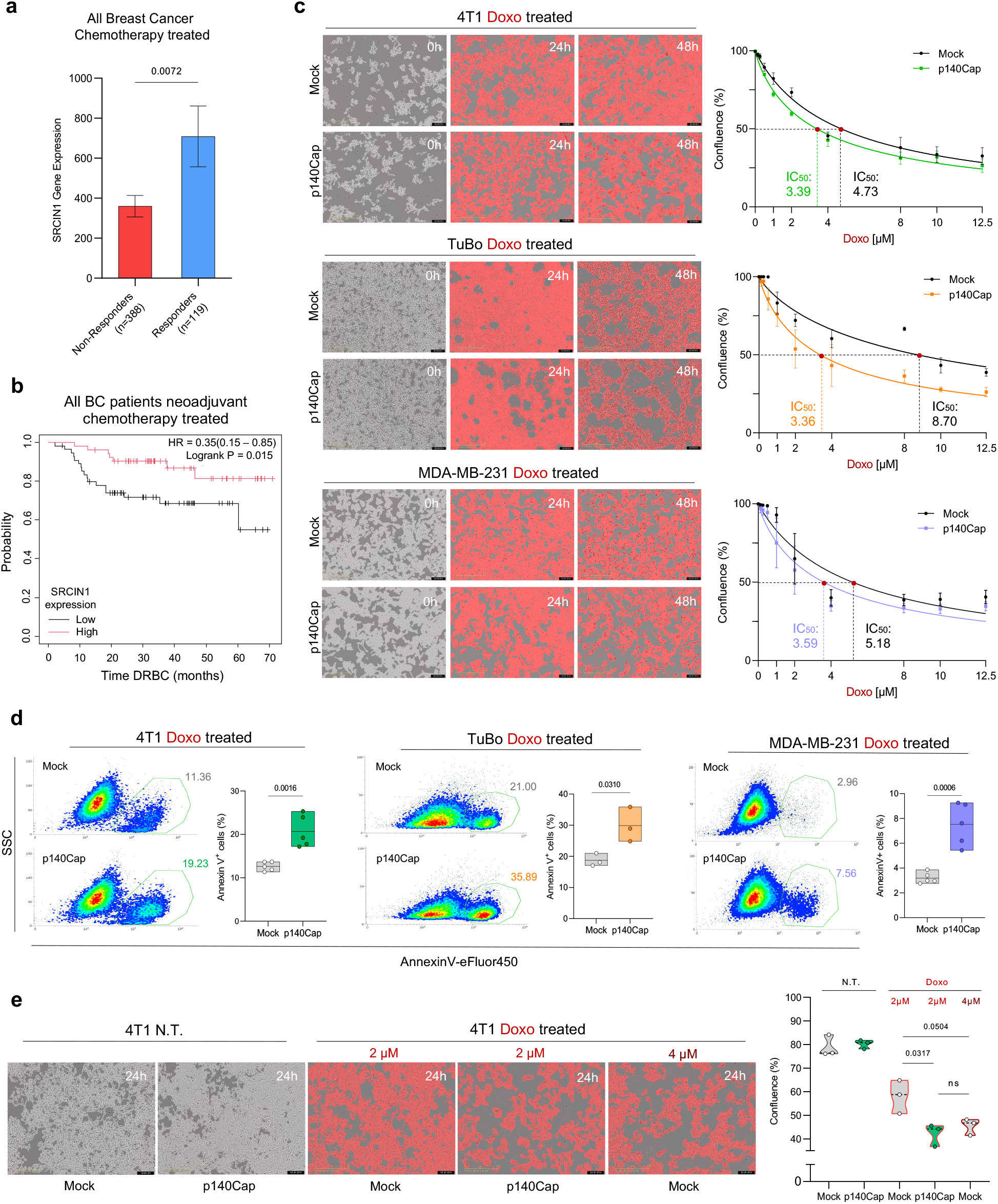
p140Cap expression enhances chemotherapy sensitivity in breast cancer. **a**, SRCIN1 expression in responders (n = 119; no residual tumour after chemotherapy) versus non-responders (n = 388; residual tumour present) in ROCplot chemotherapy-treated BC cohort^33^. Data are mean ± SEM; significance was assessed by a two-tailed unpaired t-test. **b**, Overall survival curve of BC patients treated with neoadjuvant chemotherapy, stratified by SRCIN1 median expression identified by 236838_at probe (high, *n* = 52; low, *n* = 55). Analysis was performed using an integrated clinical dataset from METABRIC, TCGA and IMPACT cohorts^34^. Time DRBC (Death Related to Breast Cancer) is reported in months. Hazard ratio (HR) with 95% confidence interval and two-sided log-rank *P* value are indicated in the plot. **c**, Representative live-cell images (left) and dose-response curves (right) of Mock and p140Cap 4T1 (images displayed: Doxo 2 μM), TuBo (images displayed: Doxo 4 μM) and MDA-MB-231 (images displayed: Doxo 2 μM) cells treated with Doxo. Cells were exposed to increasing concentrations (0-12.5 μM) of Doxo and monitored for 0-48 h with IncuCyte SX5. Quantitative analysis was performed using the Cell-by-Cell analysis module based on calculated area of cell mask; analysis masks are shown in grey (pre-treatment) and in red (post-treatment). IC50 curves generated from dose-response measurements of cell confluence 48 h after treatment. **d,** Flow cytometry plots and floating bar (min-to-max) quantification of Annexin V-positive apoptotic cells in Mock and p140Cap 4T1 (*n* = 5), TuBo (*n* = 3) and MDA-MB-231 (*n* = 5) cells after 24 h treatment with Doxo. Apoptotic cells were identified by Annexin V-eFluor450 staining and quantified by flow cytometry. **e**, Representative Incucyte images (left) and confluence truncated violin plot (right) of Mock and p140Cap 4T1 cells after 24h treatment with Doxo 2 or 4μM. Quantitative analysis was performed using the Cell-by-Cell analysis module; analysis masks are shown in grey (pre-treatment) and in red (post-treatment). Points represent biological replicates. Statistical significance was assessed using a two-tailed unpaired *t*-test.

### p140Cap improves intracellular doxorubicin retention

To delineate the doxorubicin dynamics underlying the enhanced chemosensitivity of p140Cap-expressing cells, we examined the intracellular accumulation and efflux kinetics of the drug. Taking advantage of the intrinsic red fluorescence of doxorubicin, we performed flow cytometry following low-dose treatment for 24 hours to avoid cell death and detachment. Across all three cell lines, p140Cap cells displayed a higher percentage of doxorubicin-positive cells than controls, suggesting an increased intracellular drug accumulation (Fig. 2a). Moreover, doxorubicin positivity decreased more rapidly over time in Mock MDA-MB-231 and TuBo cells than in p140Cap counterpart, consistent with sustained intracellular drug retention in p140Cap presence (Extended Data Fig. 2a,b). Doxorubicin intracellular availability, and therefore its therapeutic efficacy, is regulated by a dynamic interplay between passive diffusion across the plasma membrane and active efflux mediated by ATP-dependent transporters^12,34^. To investigate the efflux process, we treated TuBo cells with doxorubicin, followed by media washout after 6 hours, in order to remove doxorubicin excess and avoid drug re-uptake, tracking doxorubicin kinetics by IncuCyte live-cell imaging (Fig. 2b). Red counts fluorescence analysis, which reflects the amount of intracellular doxorubicin, revealed that control cells exhibit faster doxorubicin extrusion than p140Cap cells, supportive of an increased efflux (Fig. 2c,d and Supplementary Videos 1 and 2). Concomitantly, after drug removal, Mock cells reached higher confluence than p140Cap cells (Fig. 2c,e). Together, these data indicate more efficient intracellular doxorubicin clearance in Mock cells, likely via active efflux, which in turn reduces the drug’s cytotoxic efficacy. Conversely, p140Cap cells exhibit a more sensitive phenotype, with prolonged intracellular doxorubicin retention.

**Fig. 2:**
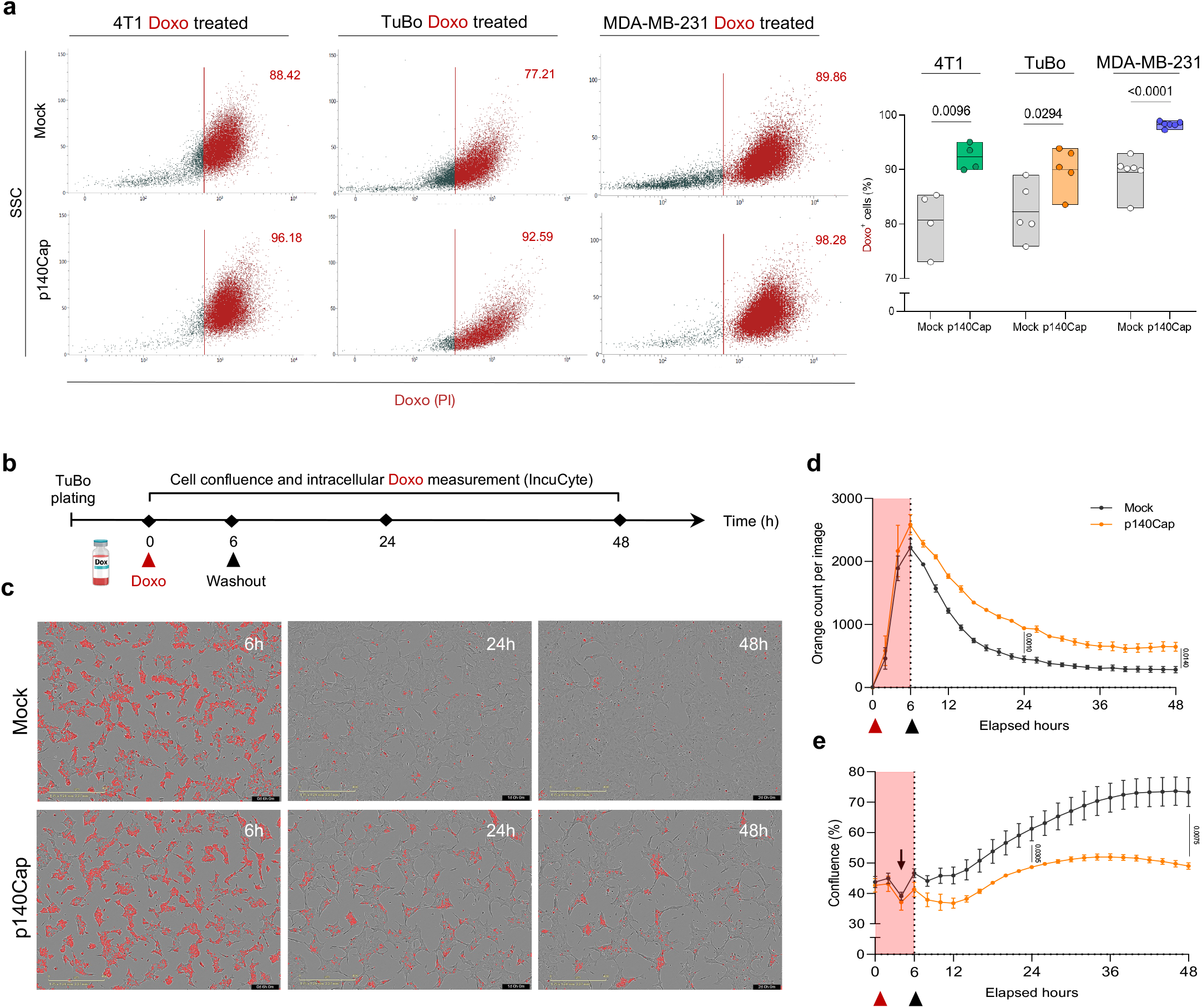
p140Cap improves intracellular accumulation of doxorubicin. **a**, Representative flow cytometry plots and floating bar (min-to-max) quantification of Doxo-positive cells in Mock and p140Cap 4T1 (*n* = 4), TuBo (*n* = 5) and MDA-MB-231 (*n* = 6) cells. Cells were treated with 0.5 µM, 0.125 µM and 0.5 µM Doxo, respectively, and analyzed 24 h post-treatment; gating was established based on the corresponding untreated control. **b**, Schematic representation of experimental set-up for Doxo efflux experiments, created with Biorender.com. **c-e**, Representative live-cell images (c) and quantitative time-course analysis of intracellular Doxo accumulation (d) and cell confluence (e) of TuBo cells. Cells were exposed to 1 μM Doxo for 6 h (red shading), followed by drug washout (black triangle), and monitored using an IncuCyte live-cell imaging system. Intracellular Doxo levels were quantified as the number of orange fluorescent objects per image, while cell confluence was assessed using integrated AI-based image analysis of IncuCyte software. Doxo accumulation was quantified every 2 h for 48 h following treatment initiation (red triangle) and washout (black triangle) by automated orange object count implemented in Incucyte software. Data are presented as mean ± SEM from three independent biological replicates. Statistical significance using unpaired two-tailed Student’s t-tests.

### ABCC1 reduction in p140Cap cells promotes chemosensitivity via increased intracellular doxorubicin retention

The observed differences in intracellular doxorubicin accumulation and chemo-responsiveness between Mock and p140Cap cells prompted us to examine the expression of the multidrug-resistance ABC transporters ABCC1, ABCG2, and ABCB1^12,13^. Interestingly, qRT-PCR profiling in 4T1, TuBo and MDA-MB-231 cells revealed a consistent selective reduction of ABCC1 levels in p140Cap-expressing BC cell models relative to controls (Fig.3a and Extended Data Fig. 3a). Concordantly, transcriptomic profiling of BC cohort from GEPIA3^38^ identified ABCC1 as the most highly expressed multidrug-resistance ABC transporter, thereby underscoring its biological and clinical relevance. (Extended Data Fig. 3b). In accordance, DepMap.org mass spectrometry-based proteomics data revealed an inverse correlation between p140Cap and ABCC1 protein levels in BC cell lines, consistent with reduced drug efflux capacity of p140Cap cells (Fig. 3b). Noteworthy, ABCC1 protein levels exhibit a positive correlation to mRNA levels in both DepMap All and BC cell lines (Extended Data Fig. 3c,d). To functionally link p140Cap chemo-sensitizing effect to reduced ABCC1 expression, we next treated TuBo and MDA-MB-231 Mock cells with reversan^39^, a pharmacological inhibitor of ABCC1. The number of doxorubicin-positive cells significantly increases following the addition of reversan, indicating that blocking ABCC1 mimics p140Cap expression (Fig. 3c,d). Moreover, co-treatment with doxorubicin and reversan reduced Mock cell viability to levels comparable to those observed in p140Cap cells treated with doxorubicin alone (Fig. 3e,f). In line with this, siRNA-mediated knockdown of ABCC1 in MDA-MB-231 Mock cells decreased cell viability following doxorubicin treatment and increased intracellular doxorubicin accumulation, closely recapitulating the sensitive phenotype observed in p140Cap cells (Extended Data Fig. 3e-g). Consistently, across independent BC cohorts from both METABRIC (*n* = 1904)^32^ and TCGA (*n* = 956)^40^ dataset, ABCC1 transcript levels are positively correlated with GRAESSMANN_RESPONSE_TO_MC_AND_DOXORUBICIN_DN and GRAESSMANN_APOPTOSIS_BY_DOXORUBICIN_DN signatures^41^, which contain genes transcriptionally repressed upon doxorubicin treatment and during doxorubicin-induced apoptosis, respectively (Fig. 3h). Overall, these results identify ABCC1 reduction as a key mediator whereby p140Cap promotes intracellular doxorubicin retention and enhances chemotherapeutic efficacy.

**Fig. 3:**
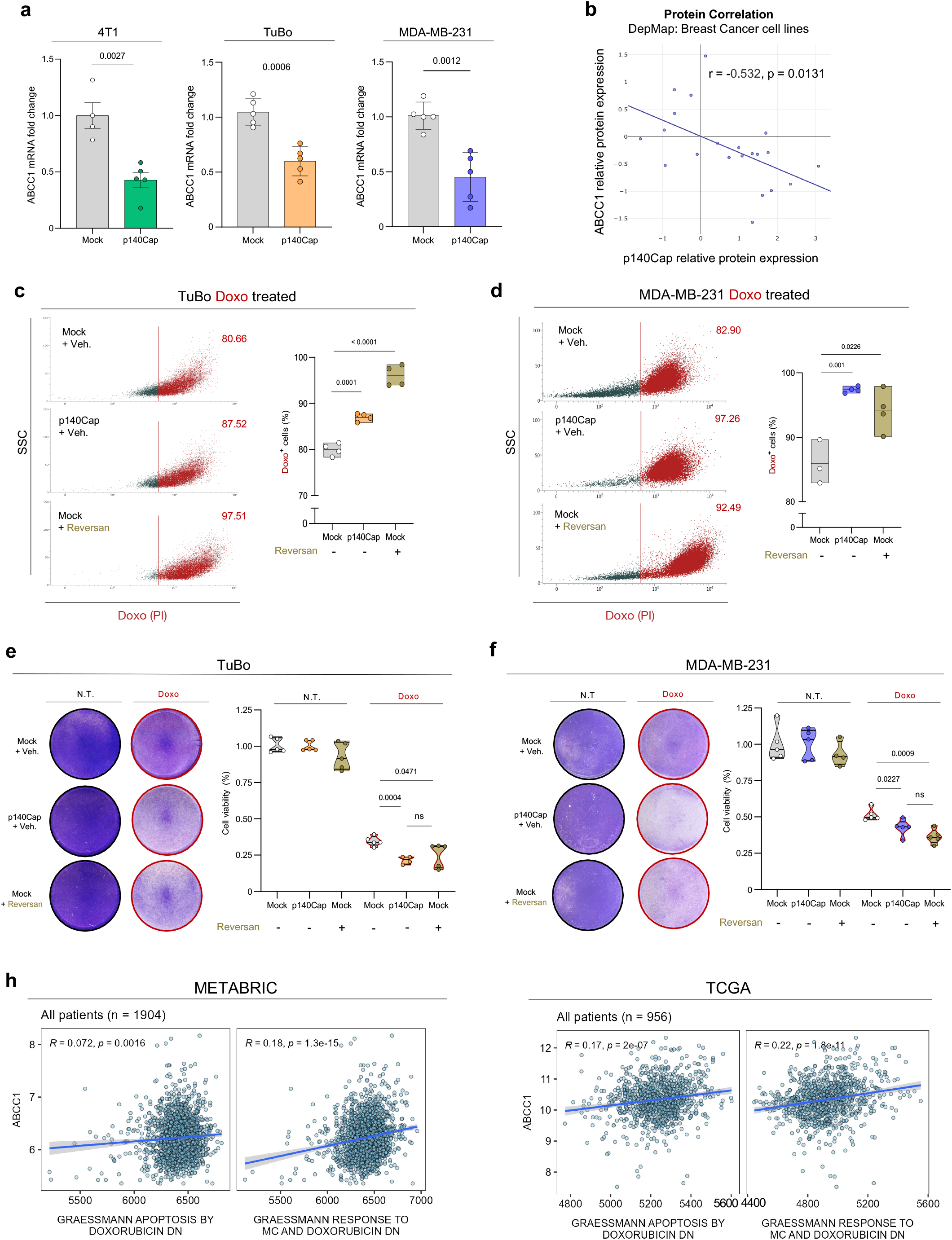
ABCC1 downregulation in p140Cap cells is associated with increased intracellular doxorubicin retention. **a,** qRT-PCR analysis of ABCC1 mRNA expression in Mock and p140Cap 4T1 (*n* = 4), TuBo (*n* = 5), and MDA-MB-231 (*n* = 5) cells. Expression levels were normalized to Mock control. Data are represented for n = x experimental repeats and shown as mean ± SEM; two-tailed unpaired t test. **b**, Correlation analysis between p140Cap (sp|Q9C0H9-5|) and ABCC1 (sp|P33527|) expression at protein levels across breast cancer cell lines from DepMap dataset. Pearson correlation analysis is indicated. **c, d**, Representative flow cytometry plots and floating bar (min-to-max) quantification of Doxo-positive cells in Mock and p140Cap TuBo (*n* = 4) and MDA-MB-231 (*n* = 4) cells. Where indicated, Mock cells were pre-treated with Reversan (10 µM) for 1 h before exposure to 1 µM Doxo. Cells were analyzed 24 h post-treatment by flow cytometry and gating was established based on the corresponding untreated control. **e**, **f**, Representative images and quantification of Crystal Violet of TuBo (e) and MDA-MB-231 (f) following 1 µM Doxo treatment. Where indicated, Mock cells were pretreated with Reversan (10 µM). Data derived from five independent experiments. Statistical analysis was performed using unpaired t-tests. **h**, Correlation between ABCC1 and the indicated gene sets in breast cancer patients in the Metabric (*n* = 1,904 patients) and TCGA (*n* = 956 patients) datasets.

### p140Cap limits an ABCC1-enriched breast cancer stem cell side population via β-Catenin inhibition

Given that the abundance of BCSCs limits chemotherapy efficacy, we reasoned that the shrinkage of Doxorubicin-Negative Side Population (DNSP; cells with undetectable intracellular doxorubicin after treatment) observed in p140Cap cells reflects a smaller BCSC compartment, likely enriched for ABCC1 expression (Fig. 2a). To prove this scenario, we isolated DNSP from doxorubicin-treated 4T1 and MDA-MB-231 Mock cells by FACS to evaluate its self-renewal potential and ABCC1 levels relative to bulk populations (Fig. 4a). The DNSP displays enhanced mammospheres-forming potential than both bulk Mock and p140Cap, consistent with BCSC enrichment (Fig. 4b). Interestingly, digital droplet PCR (ddPCR) shows that DNSP exhibits increased expression of ABCC1 transporter relative to bulk counterparts (Fig. 4c). Taken together, these findings highlight that p140Cap presence markedly reduces the ABCC1-enriched DNSP, a key reservoir of chemotherapy-unresponsive BCSCs. Concordantly, (i) in our models the ABCC1 expression is significantly higher in BCSC-enriched mammospheres compared to 2D cell cultures (Extended Data Fig. 4a); (ii) from both TNBC and HER2 TCGA BC cohort emerges a positive and negative correlation between ABCC1 expression and CONRAD STEM CELL and LIM MAMMARY STEM CELL DOWN^42^ genes signatures, respectively (Extended Data Fig. 4b); (iii) from DepMap database, reveal a positive correlation between ABCC1 expression and MALTA_CURATED_STEMNESS_MARKERS^43^, LIM_MAMMARY_STEM_CELL_UP^42^ and WONG_ADULT_TISSUE_STEM_MODULE^44^ signatures (Extended Data Fig. 4c,d). p140Cap is known to reduce the active β-Catenin pool by stabilizing the β-Catenin destruction complex^29^. Considering the critical role of this pathway in maintaining stemness, we investigated whether modulation of β-Catenin signaling could influence ABCC1 expression and thereby affect chemosensitivity. To test this, we expressed in TuBo p140Cap cells a constitutively active (C.A.) form of β-Catenin (S33Y) that cannot be phosphorylated, ubiquitinated and subsequently degraded^45^. Importantly, the S33Y construct is able to (i) diminish doxorubicin intracellular accumulation; (ii) rescue cell viability; (iii) restore ABCC1 expression in TuBo p140Cap cells, confirming that the chemo-sensitizing effects of p140Cap critically depend on its ability to counteract β-Catenin signaling (Fig. 4d-f). Collectively, these data identify p140Cap as a determinant of doxorubicin responsiveness, acting through β-Catenin signaling to limit an ABCC1-enriched BCSC compartment and thereby enhance chemotherapy response.

**Fig. 4:**
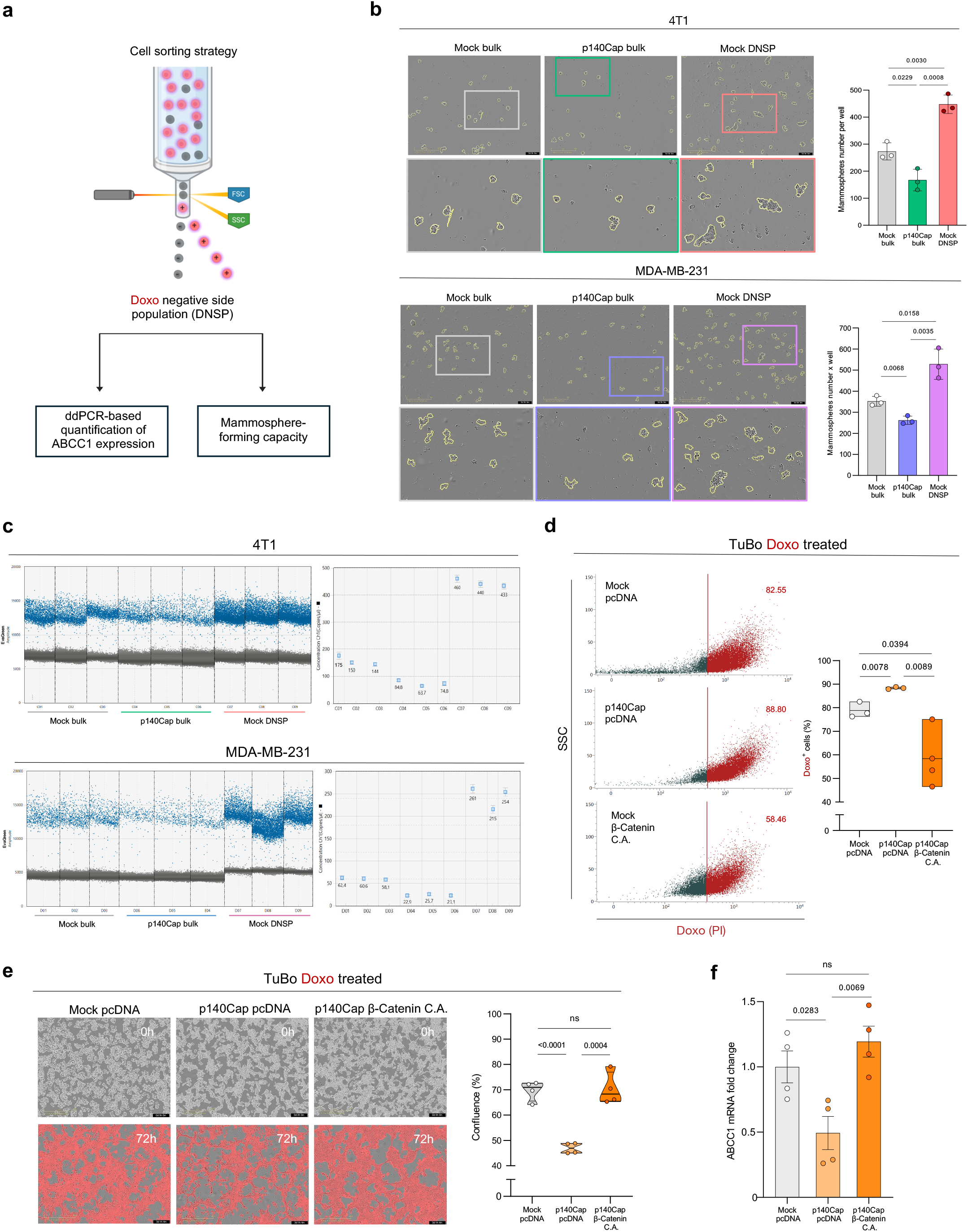
p140Cap limits an ABCC1-enriched breast cancer stem cell side population via β-Catenin inhibition. **a,** Flow cytometric identification and sorting strategy of the doxorubicin-negative side population (DNSP). Black points indicate Doxo-negative cells used for ddPCR quantification of ABCC1 expression and mammospheres-forming potential. Created with BioRender.com. **b**, Representative images and quantification of mammospheres formation in Mock and p140Cap 4T1 (*n* = 3) and MDA-MB-231 (*n* = 3) cells cultured under low-attachment conditions. Data show the number of spheres per well. **c**, ddPCR-based absolute quantification of ABCC1 in 4T1 and MDA-MB-231 sorted cells. (Left) 1D EvaGreen amplitude plot showing ABCC1 expression in Mock bulk (grey), p140Cap bulk (green) and Mock DNSP (red). (Right) ABCC1 (Ch1) copy number per µL derived from QX Manager Standard Edition (2.2.0). **d**, Representative flow cytometry plots and floating bar (min-to-max) quantification of Doxo-positive cells in TuBo Mock pcDNA (*n* = 3), p140Cap pcDNA (*n* = 3) and p140Cap β-Catenin C.A. (*n* = 4) cells. Cells were treated with Doxo 0.125 µM, and analyzed 24 h post-treatment; gating was established based on the corresponding untreated control. **e**, Representative live-cell images of TuBo cells expressing Mock pcDNA, p140Cap pcDNA, or p140Cap β-Catenin C.A. treated with 1 µM Doxo and imaged 72 h post-treatment using the IncuCyte SX5 system. Cell-by-Cell analysis masks are shown in grey (pre-treatment) and red (post-Doxo) to quantify cell confluence. Data represent *n* = 4 biological replicates. **f**, qRT-PCR analysis of ABCC1 mRNA in TuBo cells expressing Mock pcDNA, p140Cap pcDNA, and p140Cap β-Catenin C.A.. Expression levels were normalized to Mock controls. Data represent *n* = 4 biological replicates and are shown as mean ± SEM. Statistical significance was determined by two-tailed unpaired *t* test.

### Pharmacological inhibition of β-Catenin signaling restores chemosensitivity in p140Cap-deficient breast cancer cells

Pharmacological strategies that mimic p140Cap role may provide a rational and selective approach to target β-Catenin and DNSP to improve chemo efficacy in p140Cap-negative models. As proof of concept, we pre-treated TuBo and MDA-MB-231 Mock cells with two doses of IWR-1, an AXIN1 pharmacological destruction complex stabilizer able to constrain the BCSC compartment through induction of β-Catenin degradation (Fig. 5a)^46^. As expected, pretreatment with IWR-1 reduced Sca-1^47^ expression, a putative BCSC marker (Extended Data Fig. 5a). Interestingly, in Mock cells IWR-1 pre-treatment followed by doxorubicin administration: (i) increases intracellular drug retention; (ii) reduces cell viability; (iii) increase doxorubicin-dependent apoptosis; (iiii) suppresses ABCC1 expression, closely mirroring the p140Cap chemosensitive phenotype (Fig. 5b-e). To directly assess the impact on the BCSCs, we generated TuBo-derived mammospheres, pre-treated with IWR-1, dissociated into single cells, and subsequently treated with doxorubicin. Notably, IWR-1 pretreatment reduced the BCSC pool, as assessed by Sca-1 expression, and increased intracellular doxorubicin retention to p140Cap-derived mammospheres levels (Extended Data Fig. 5b,c). To extend these findings toward clinical relevance, we evaluated LGK-974, a small-molecule inhibitor of Wnt signaling that has completed phase I clinical trial in TNBC patients^48,49^. LGK-974 blocks PORCN-dependent Wnt secretion, thereby suppressing Wnt/β-Catenin signaling and reducing the BCSC compartment. Similar to IWR-1, when administered as a pre-treatment (Fig. 5f), LGK-974 restored doxorubicin sensitivity in TuBo and MDA-MB-231 Mock cells, recapitulating the chemo-responsive phenotype conferred by p140Cap expression (Fig. 5g). Together, these findings provide functional evidence that pharmacological strategies aimed at shrinking the BCSC compartment via β-Catenin pathway inhibition phenocopy p140Cap biology and enhance doxorubicin sensitivity in BC cells. Accordingly, targeting β-Catenin signaling with IWR-1 or LGK-974 may represent a promising therapeutic approach to sensitize p140Cap-deficient breast tumors.

**Fig. 5:**
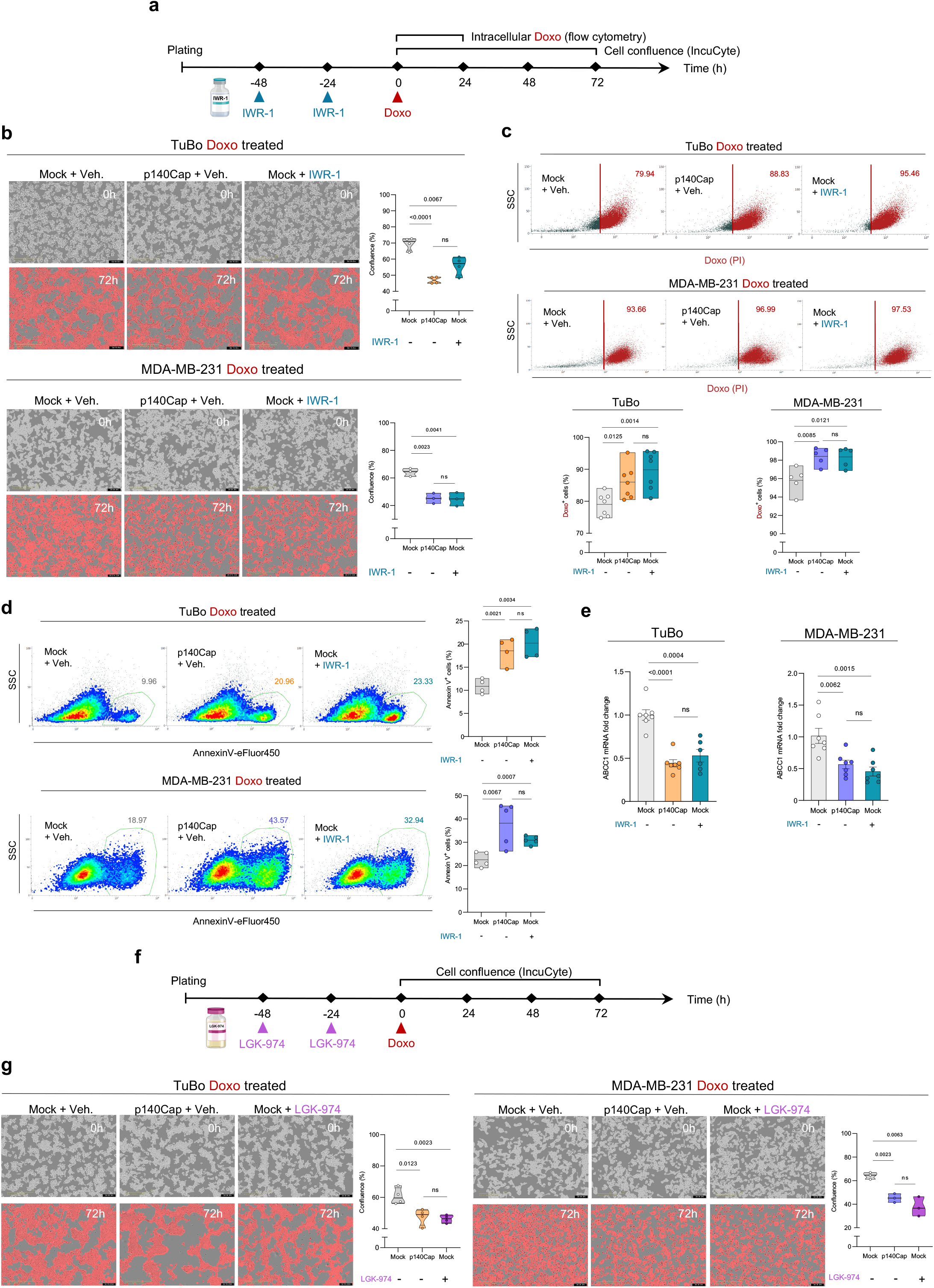
Pharmacological inhibition of β-Catenin signaling phenocopies p140Cap presence and restores chemosensitivity in breast cancer cells. **a,** Schematic representation of the treatment schedule for IWR-1 and Doxo administration. Experimental timeline and drug exposure sequence are illustrated (created with BioRender.com). **b,** Representative live-cell images and IncuCyte-based quantification of cell confluence in TuBo and MDA-MB-231 cells treated with 1 µM Doxo in the presence (+) or absence (-) of the Wnt/β-Catenin inhibitor IWR-1 (5 µM). Points represent biological replicates. **c,** Representative flow cytometry plots and quantification of intracellular Doxo fluorescence in Mock and p140Cap TuBo (*n* = 7) and MDA-MB-231 (*n* = 5) cells following 24 h Doxo treatment, with or without IWR-1 pre-treatment (+). Percentages of Doxo-positive cells are indicated in red; gating was established based on the corresponding untreated control. **d**, Representative flow cytometry plots and quantification of Annexin V^+^ apoptotic cells in Mock and p140Cap TuBo cells treated with Doxo (0.5 µM) for 72h in the presence or absence of IWR-1 (5 µM) pre-treatment (+). **e**, qRT-PCR analysis of ABCC1 mRNA expression comparing Mock, p140Cap and Mock pre-treated with IWR-1 in TuBo and MDA-MB-231 cells. Data represent mean ± SEM from *n* = 7 independent experiments. **f**, Schematic representation of the treatment schedule for LGK-974 and Doxo administration. Experimental timeline and drug exposure sequence are illustrated (created with BioRender.com). **g**, Representative live-cell images and IncuCyte-based quantification of cell confluence in TuBo and MDA-MB-231 cells treated with 1 µM Doxo in the presence or absence of the Wnt/β-Catenin inhibitor LGK-974 (5 µM). Points represent biological replicates as indicated.

### p140Cap improves doxorubicin response in preclinical models of breast cancer

To evaluate the *in vivo* relevance of p140Cap-mediated chemosensitivity, we orthotopically injected Mock and p140Cap TuBo and 4T1 cells into the left mammary fat pads of six/eight-week-old syngeneic BALB/c female mice. In TuBo tumor-bearing mice, doxorubicin was administered at 3 mg/kg every two days for seven intraperitoneal injections, a regimen consistent with standard preclinical dosing (Fig. 6a). In this setting, p140Cap-expressing tumors exhibited significantly reduced tumour growth upon doxorubicin treatment compared with controls (Fig. 6b). In parallel, 4T1 tumor-bearing mice were treated with a reduced doxorubicin dose of 1.5 mg/kg, following the same schedule, to assess whether p140Cap expression could sustain therapeutic efficacy under low-dose conditions (Fig. 6a). Under this conditions, p140Cap-expressing tumors showed markedly impaired growth and a lower number of lung metastases compared with Mock controls (Extended Data Fig. 6a,b). Consistent with *in vitro* data, p140Cap TuBo tumors displayed reduced ABCC1 expression compared to Mock tumors (Fig. 6c). As previously reported^29^, untreated TuBo and 4T1 tumors expressing Mock or p140Cap showed baseline differences in growth, which obscure differential responses to doxorubicin, hindering hypothesis validation. We therefore employed a substantially lower doxorubicin dose (0.5 mg/kg) in 4T1 tumor-bearing mice, corresponding to one-third of the previous dosage, following the treatment schedule as in Fig. 6d. Notably, this dose had no measurable effect on Mock tumors compared with untreated controls, whereas p140Cap-expressing tumors remained significantly responsive to treatment (Fig. 6e). These findings indicate that p140Cap lowers the dose threshold required for an effective antitumor response in vivo. To determine whether pharmacological inhibition of β-Catenin could reproduce the p140Cap phenotype, Mock tumor-bearing mice were treated with IWR-1 in combination with doxorubicin according to the schedule shown in Fig. 6d. Importantly, IWR-1 sensitized Mock tumors to doxorubicin treatment, recapitulating the response observed in p140Cap-expressing tumors and supporting β-Catenin inhibition as a potential strategy to restore chemosensitivity in p140Cap-deficient tumors (Fig. 6f). In line with the primary tumour growth, 4T1 Mock tumour-bearing mice displayed a higher number of lung metastases compared with p140Cap counterparts, which exhibited consistently low metastatic lesions in both untreated and doxorubicin-treated conditions. (Extended Data Fig. 6c). As a proof-of-concept of the enhanced efficacy of doxorubicin in p140Cap tumour-bearing mice, we assessed DNA damage and consequent apoptosis by quantifying γH2AX and cleaved caspase-3, respectively. Notably, low-dose doxorubicin significantly increased both markers selectively in p140Cap-expressing tumors, indicating enhanced sensitivity to chemotherapy in vivo (Fig. 6g, h). Moreover, normalization of the tumour vasculature can enhance perfusion, vascular permeability and intratumoral drug distribution^50–52^, potentially contributing to p140Cap-mediated chemo-sensitization in vivo. To determine whether p140Cap influences tumour vascular architecture, we quantified CD31⁺CD105⁺ endothelial cells and found comparable vascularization patterns in size-matched Mock and p140Cap TuBo tumours (Fig. 6i). Strikingly, p140Cap TuBo tumors showed a significant enrichment of immature NG2⁺ and mature α-SMA⁺ putative pericytes coverage markers^53^, indicative of more normalized and stabilized tumor vasculature (Fig. 6l,m). Overall, these data demonstrate that p140Cap enhances doxorubicin efficacy in vivo by reducing primary tumor growth and metastatic dissemination. In addition to its tumor cell-intrinsic effects on the ABCC1-enriched stem-like compartment, p140Cap may further improve chemotherapy response through vascular normalization, thereby facilitating drug perfusion and intratumoral distribution. Pharmacological stabilization of the β-Catenin destruction complex with IWR-1 provides an alternative strategy to improve doxorubicin response in p140Cap-deficient tumors.

**Fig. 6:**
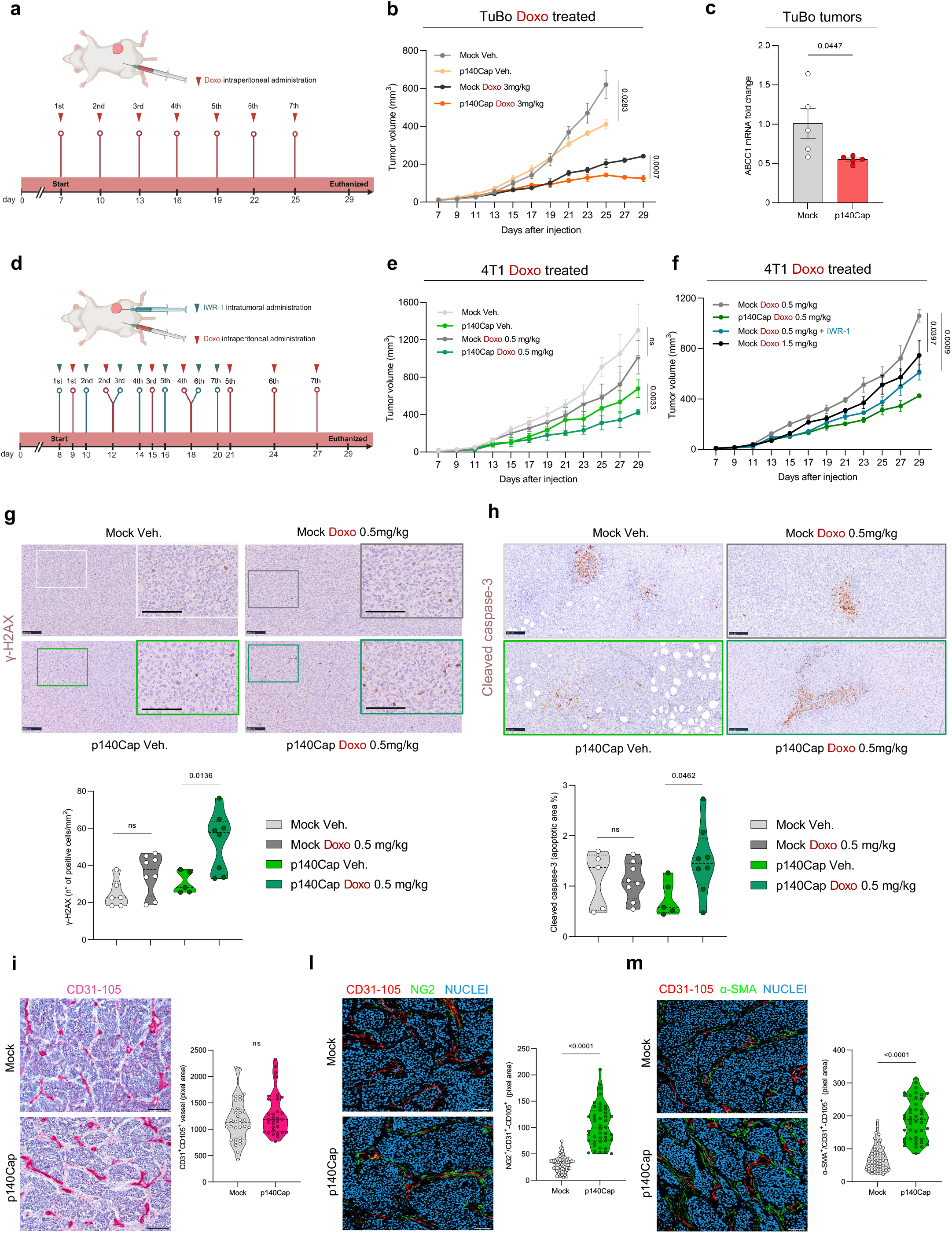
p140Cap improves doxorubicin response in preclinical models of breast cancer. **a**, Experimental scheme for the TuBo and 4T1 orthotopic model. Tumor cells were injected into the mammary fat pad and mice received seven intraperitoneal Doxo treatments according to the indicated schedule before endpoint. Created with BioRender.com **b,** Tumor growth curves of TuBo Mock and p140Cap tumors treated with vehicle (Veh.) or Doxo (3mg/kg). Tumor volume was measured with caliper every 2 days. **c**, ABCC1 mRNA fold change in Mock and p140Cap TuBo tumors, assessed by qRT-PCR. Dots represent individual mice and bars indicate mean ± SEM. **d**, Experimental scheme for the 4T1 orthotopic model. Mice received seven intraperitoneal Doxo (0.5mg/kg) and/ intratumoral IWR-1 (5mg/kg) according to the indicated dosing schedule. **e**, Tumor growth curves of 4T1 Mock or p140Cap tumors treated with Veh. or Doxo, as indicated. **f**, Tumor growth curves of 4T1 tumors treated with Doxo (0.5 mg/kg), Doxo (1.5mg/kg), or Doxo (0.5mg/kg) combined with IWR-1, as indicated. In b, e and f, data are mean ± SEM. **g,** Representative images of immunohistochemical staining for γ-H2AX showing DNA damage in 4T1 tumors. Each tumor was divided into two distinct regions, which were analyzed separately; violin plots illustrate the resulting data distributions. Scale bars, 100 μm. **h,** Representative images of immunohistochemical staining for cleaved caspase-3 performed on 4T1 tumor sections from the indicated treatment groups and quantification of the apoptotic area. **i-m**, Representative IHC and IF images of TuBo tumor vasculature. For immunohistochemistry, tumor cryosections were stained with CD31 mixed with CD105 to visualize endothelial cells (i). For immunofluorescence, cryosections were stained for CD31 and CD105 (endothelial cells), NG2 and α-SMA (pericytes); nuclei were counterstained with DRAQ5. Exact *P* values are shown in the panels; Statistical analyses were performed using two-tailed unpaired *t*-tests.

### p140Cap is a candidate predictive biomarker of chemotherapy response in breast cancer

To assess the clinical relevance of p140Cap, we focused on TNBC, the most aggressive breast cancer subtype, in which chemotherapy remains a cornerstone of treatment. Proteomic analysis of a TNBC cohort (DLDCCC)^54^ revealed an inverse correlation between p140Cap and ABCC1 protein expression (Fig. 7a). To further investigate this observation, we selected three TNBC PDX models displaying different endogenous levels of p140Cap expression, including two p140Cap-low/negative models (Pt A and Pt B) and one p140Cap-high model (Pt C), as confirmed by immunohistochemistry (Fig. 7b). Notably, ABCC1 mRNA levels were significantly lower in Pt C than in Pt A and Pt B (Fig. 7c), supporting an inverse association between p140Cap and ABCC1 expression in clinically relevant tumor models. To directly investigate the functional relevance, p140Cap was re-expressed in the p140Cap-low PDX models (Pt A and Pt B) using a lentiviral construct, as confirmed by immunofluorescence (Fig. 7d). Importantly, p140Cap re-expression consistently reduced ABCC1 expression (Fig. 7e), increased intracellular doxorubicin accumulation (Fig. 7f) and enhanced doxorubicin sensitivity (Fig. 7g), indicating that restoration of p140Cap is sufficient to promote a chemo-sensitive phenotype in patient-derived tumor cells. To further support these results at the single-cell level, analysis of scRNA-seq data from two TNBC patients^55^ showed a trend toward mutually exclusive expression of *SRCIN1* and *ABCC1* within distinct epithelial tumor cell subpopulations (Fig. 7h). Collectively, these findings establish a functional link between p140Cap, ABCC1 expression and doxorubicin responsiveness in patient-derived TNBC models. They support p140Cap as a candidate predictive biomarker of chemotherapy response and identify p140Cap-deficient tumours as a subgroup potentially amenable to combinatorial therapeutic strategies aimed at restoring chemosensitivity.

**Fig. 7:**
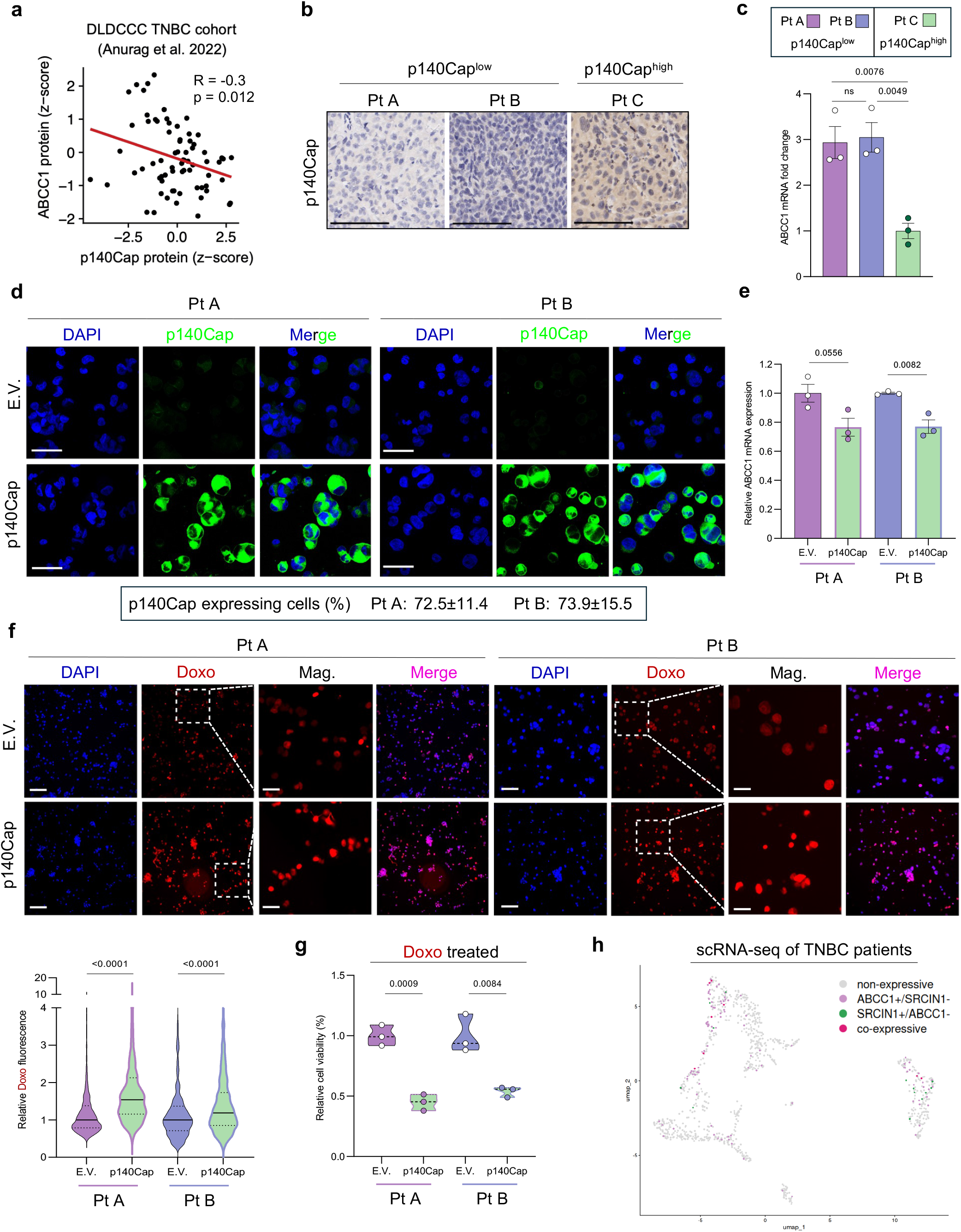
p140Cap as a positive predictive biomarker of chemotherapy efficacy in breast cancer patients. **a,** Inverse correlation between p140Cap (SRCIN1) and ABCC1 protein expression levels in TNBC from the DLDCCC cohort. R, Pearson’s correlation coefficient; p, Pearson’s correlation test; *n* = 72. **b,** Representative IHC staining of p140Cap, p140Cap^Low^ (Pt A and Pt B) and p140Cap^High^ (Pt C) PDX models. Bar, 100 μm. **c**, qRT-PCR analysis of ABCC1 mRNA expression in the PDX models. Expression levels were normalized to the p140Cap^High^ (Pt C). Data represent *n* = 3 biological replicates and are shown as mean ±SEM. **d,** Representative fluorescence images of PDX cells lentivirally transduced with p140Cap or empty vector (E.V.) and co-stained for p140Cap and DAPI, Bars, 50 μm. Box shows quantification of p140Cap-expressing cells. Data are shown as the percentage of p140Cap-positive cells ± SD. **e,** qRT-PCR analysis of ABCC1 mRNA expression in E.V. and p140Cap PDX cells. Expression levels were normalized to the E.V. condition. Data represent *n* = 3 biological replicates and are shown as mean ± SEM. **f,** Representative fluorescence images of PDX cells incubated with 1 μM Doxo for 1 h and stained with DAPI. Bar, 100 μm, Mag. 25 μm. Violin plot with quantification of nuclear doxorubicin fluorescence intensity. Fluorescence values were normalized to the median intensity of the corresponding control E.V. sample and are presented as relative mean doxorubicin fluorescence. Data are shown as median ± SEM. **g,** 3D *in vitro* growth of PDX cancer cells stably transduced with either p140Cap or the E.V. as control and treated with Doxo 20 nM, was evaluated by direct cell counting. Data are expressed relative to Time 0 for each condition and represent the mean ± SEM (*n* = 3). Statistical significance was determined using a two-tailed unpaired t-test. **h**, UMAP of SRCIN1 and ABCC1 expression in basal-like TNBC cells. Basal-like epithelial cells (*n* = 1,013) isolated from scRNA-seq data of two TNBC patients (GSE263995) were classified as in figure.

## Discussion

Our findings identify p140Cap as a determinant of BC responsiveness to chemotherapy, linking its expression to an increased chemosensitivity across murine, human and patient-derived BC models. Notably, p140Cap is expressed in approximately 50% of cases in both HER2-positive and TNBC^26,29^, underscoring its broad prevalence across distinct clinically relevant molecular subtypes beyond a single disease context. The consistent enhancement of doxorubicin sensitivity observed in p140Cap-expressing tumors supports a functional role for p140Cap in shaping therapeutic response and suggests that p140Cap-positive tumors may achieve clinically meaningful benefit under reduced chemotherapy dosage. In this context, p140Cap-mediated chemo-sensitization raises the possibility of moving beyond conventional maximum tolerated dose regimens toward low-dose metronomic strategies, which could preserve antitumor efficacy while limiting systemic toxicity^56^. Such an approach may be particularly relevant for anthracycline-based therapies, whose cumulative adverse effects frequently restrict long-term administration^34,57^. Beyond improving tolerability and patient quality of life, reducing the need for high-dose chemotherapy could also alleviate the substantial economic burden associated with treatment-related toxicities, supportive care and hospitalization^58^. Together, these findings position p140Cap not only as a candidate biomarker of chemotherapy responsiveness, but also as a potential biological framework for the development of more sustainable and less toxic therapeutic strategies in BC (Fig. 8).

**Fig. 8:**
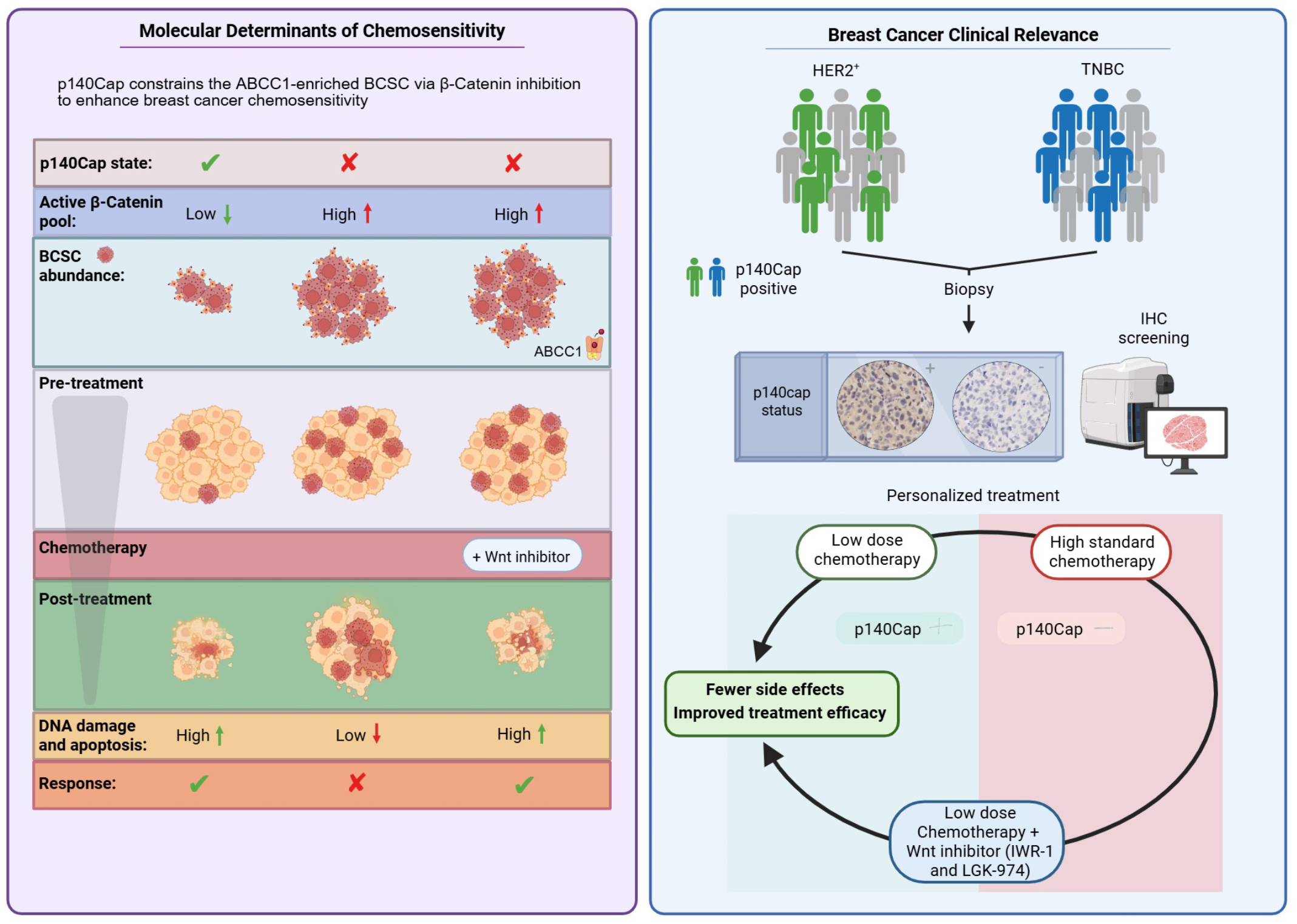
p140Cap constrains an ABCC1-enriched breast cancer stem cell compartment through β-Catenin inhibition and supports therapeutic stratification in breast cancer. Left, schematic model illustrating how p140Cap regulates chemotherapy response through modulation of β-Catenin signaling and BCSC abundance. In p140Cap-positive tumors, impaired β-Catenin pathway limits the expansion of an ABCC1-enriched BCSC compartment, resulting in increased intracellular chemotherapy retention and enhanced chemotherapy-induced DNA damage and apoptosis. By contrast, p140Cap-deficient tumors display increased β-Catenin active pool, expansion of ABCC1-high BCSCs and reduced chemosensitivity. Pharmacological inhibition of Wnt/β-Catenin signaling restores chemo responsiveness in p140Cap-deficient settings, reducing residual tumour burden after treatment. Right, proposed translational framework for p140Cap-guided therapeutic stratification in HER2-positive and TNBC. Assessment of p140Cap expression by immunohistochemistry (IHC) may support treatment selection and optimization. p140Cap-positive tumors, characterized by increased intrinsic chemosensitivity, may benefit from reduced-dose chemotherapy while maintaining therapeutic efficacy and limiting toxicity. Conversely, p140Cap-negative tumors may require standard chemotherapy regimens or combination approaches incorporating Wnt/β-Catenin inhibitors, including IWR-1 or LGK-974, to restore drug sensitivity and improve treatment response. Created with BioRender.com.

A key mechanistic insight emerging from the current work is that p140Cap restrains intracellular drug clearance. Mock cells displayed more efficient doxorubicin efflux, consistent with active transport mechanisms that limit cytotoxic exposure, whereas p140Cap-expressing cells retained the drug for longer periods, thereby sustaining intracellular doxorubicin availability and enhancing therapeutic efficacy. In this context, ABCC1 reduction emerged as a critical mediator of the p140Cap-sensitive phenotype, linking p140Cap expression to impaired drug efflux and increased chemosensitivity. Although ABCC1 is a well-established driver of multidrug resistance^13,58^, direct pharmacological inhibition of this transporter has shown limited clinical success because of unacceptable systemic toxicities, likely reflecting its essential physiological functions in normal tissues^12,13,59,60^. The present study therefore do not support ABCC1 itself as a therapeutic target, but rather suggest that modulation of upstream pathways controlling ABCC1 expression and stemness-associated programs may represent a more feasible strategy to improve chemotherapy outcomes.

Beyond drug transport per se, our data indicate that p140Cap alters the composition of the chemotherapy-unresponsive compartment within breast tumors. Specifically, p140Cap markedly reduced an ABCC1-enriched doxorubicin-negative side population (DNSP) enriched for BCSCs. Unlike conventional side population assays based on fluorescent dye exclusion^61,62^, this compartment was functionally identified through doxorubicin retention itself, thereby directly linking drug efflux activity to chemotherapy sensitivity. To our knowledge, these findings describe a previously unrecognized doxorubicin-defined side population in BC associated with stemness features and reduced chemotherapy sensitivity. This is particularly relevant because stem cell-like populations are often implicated in relapse, metastatic competence and therapy evasion^11,63^. In this context, the abundance of the DNSP could itself represent a negative predictive feature of chemotherapy responsiveness, reflecting the presence of a tumor subpopulation intrinsically less sensitive to cytotoxic exposure. Future molecular and functional characterization of this compartment will be important to determine its potential translational and clinical relevance. Thus, while chemotherapy response is inherently multifactorial, our findings identify restriction of the ABCC1-enriched BCSC compartment as a major mechanism underlying the chemo-sensitizing effects of p140Cap. Nevertheless, additional tumour cell-intrinsic and microenvironmental processes are likely to contribute to the overall therapeutic response associated with p140Cap expression. In this regard, our in vivo data indicate that p140Cap tumors exhibit a more normalized vasculature, a feature that may enhance drug perfusion and intratumoral distribution, thereby providing an additional mechanism to improve chemotherapy efficacy^50–52^.

Mechanistically, these effects converge on β-Catenin signaling. The finding that pharmacological targeting of this pathway phenocopied p140Cap, reducing the stem-like compartment and enhancing doxorubicin sensitivity, provides functional support for a model in which p140Cap promotes chemosensitivity by limiting β-Catenin-dependent programs^29^. In accordance, expression of a constitutively active β-Catenin mutant abolished the p140Cap-associated phenotype, restoring ABCC1 expression, reducing intracellular doxorubicin retention and rescuing cell viability. These findings strengthen the functional link between p140Cap, β-Catenin inhibition and suppression of the stem-like compartment^29^. At the same time, they raise the possibility that β-Catenin signaling may contribute more directly to ABCC1 regulation, potentially through transcriptional regulation of transporter expression, rather than exclusively through expansion of an ABCC1-high cancer stem cell population, a hypothesis that warrants further investigation. This is conceptually important because it places p140Cap as a negative regulator of a pathway with well-established roles in stemness, poor drug responses and tumor progression. In this context, targeting β-Catenin signaling may represent a rational strategy to recapitulate p140Cap function in p140Cap-deficient tumors^17,18^. Notably, treatment with IWR-1^46^, through stabilization of the β-Catenin destruction complex, was sufficient to reproduce key features of the p140Cap-associated phenotype, supporting further investigation of this class of compounds in translational and potentially clinical settings. From a translational standpoint, the druggability of the Wnt/β-Catenin axis is supported by several therapeutic strategies currently under investigation, including the porcupine inhibitor LGK-974 in early-phase clinical trials in advanced BC^48,49^. By blocking PORCN-dependent palmitoylation and secretion of Wnt ligands, LGK-974 suppresses upstream activation of Wnt/β-Catenin signaling, thereby limiting stemness-associated programs and tumour cell plasticity^48^. Additional approaches, such as emerging targeting upstream Wnt ligand-receptor interactions and repositioned compounds like pyrvinium pamoate, which promotes β-Catenin proteasomal degradation, further highlight alternative means to attenuate this signaling axis at different regulatory levels. In this context, the development of PROTAC-based strategies targeting β-Catenin is also gaining interest as a potential next-generation approach for selective pathway modulation. Collectively, these strategies underscore the therapeutic applicability of targeting Wnt/β-Catenin signaling, although their clinical efficacy remains to be fully established^14,17,19,20,64^. Together, these observations strengthen the rationale for therapeutically targeting Wnt/β-Catenin signaling to improve chemotherapy efficacy and reducing systemic toxicity in p140Cap-deficient breast tumors (Fig. 8).

The translational relevance of our *in vitro* and *in vivo* data are reinforced by their validation in BC patient’s cohorts and patient-derived xenograft (PDX) models, which recapitulate the differential response associated with p140Cap expression. These findings support p140Cap as a candidate predictive biomarker within a precision medicine framework, in which tumors could be stratified according to p140Cap status to better anticipate responsiveness to anthracycline-based chemotherapy. In this context, p140Cap-positive tumors appear intrinsically more chemo-sensitive, raising the possibility of achieving effective disease control with reduced drug exposure and improved therapeutic index, whereas p140Cap-deficient tumors may require combination strategies aimed at restoring drug sensitivity (Fig. 8). Such stratification could inform treatment selection and dose optimization, enabling a more biologically guided therapeutic approach while reducing treatment-related toxicity and the economic burden associated with ineffective interventions. More broadly, our data position p140Cap not only as a biomarker of response, but also as a functional node whose downstream pathways may be therapeutically exploited to improve the balance between efficacy and tolerability in BC treatment.

## Methods

### Ethical approvals

All syngeneic mouse experiments and PDX studies were conducted in compliance with institutional and national guidelines for the care and use of laboratory animals. Experimental protocols were reviewed and approved by the Italian Ministry of Health (protocol no. CC652.211). All procedures were designed to minimize animal suffering and to reduce the number of animals used, in accordance with the principles of the 3Rs (Replacement, Reduction and Refinement). Research involving the collection and use of human clinical specimens was approved by the Institutional Ethics Committee/Institutional Review Board of the European Institute of Oncology (IEO, Milan, Italy), and all samples were collected under standard operating procedures with written informed consent obtained from all participants prior to collection, in accordance with approved ethical guidelines.

### Cell lines

Mycoplasma-free TuBo cells, originally derived from a spontaneous mammary carcinoma arising in BALB/c-MMTV-NeuT mice^33^, were maintained in DMEM supplemented with 20% fetal bovine serum (FBS). The murine BC cell line 4T1 (ATCC CRL-2539) and the human TNBC cell line MDA-MB-231 (ATCC HTB-26) were obtained from ATCC (LGC Standards S.r.l., Italy), authenticated according to the supplier’s recommendations and routinely tested negative for mycoplasma contamination. 4T1 cells were cultured in RPMI-1640 supplemented with 10% FBS, whereas MDA-MB-231 cells were maintained in DMEM supplemented with 10% FBS. All culture media and supplements, including FBS and penicillin-streptomycin (1%), were purchased from Invitrogen (Carlsbad, CA, USA).

### Retrovirus production and cell infection

To generate stable p140Cap-expressing cell lines, the full-length p140Cap cDNA was cloned into the pBabe-puro retroviral vector and used for retrovirus production in Platinum Retroviral Packaging cells (Cell Biolabs). Packaging cells were transiently transfected in 6-cm dishes using Lipofectamine 2000 (Thermo Fisher Scientific), according to the manufacturer’s instructions. Viral supernatants were collected 48 h post-transfection and applied directly to subconfluent TuBo, 4T1 and MDA-MB-231 target cells seeded in 6-well plates. 48 hours after infection, cells were washed and subjected to puromycin selection (1 mg/ml; Sigma-Aldrich).

In our recent work^29^, p140Cap expression was validated by western blotting and immunofluorescence using a mouse monoclonal anti-p140Cap antibody generated at the Antibody Production Facility of the Department of Molecular Biotechnology and Health Sciences, University of Torino. Antibodies were raised against a recombinant GST-fused p140Cap fragment corresponding to amino acids 800-1000 of the murine *SRCIN1* gene. To establish stable populations, infected cells were plated at single-cell density in 96-well plates (Corning), and individual puromycin-resistant clones were expanded. After 20 days of selection, p140Cap-positive clones were screened by western blotting and immunofluorescence. To minimize clonal variability, 4 independent positive clones were pooled and used for all subsequent experiments.

### Drug treatments and cell viability assay

TuBo and MDA-MB-231 cells were seeded in 6-, 24-, 48- or 96-well plates (Corning), allowed to adhere overnight, and subsequently treated with the indicated compounds under standard culture conditions (37 °C, 5% CO2). Doxorubicin (Sigma, Cat#D1515) and vinorelbine (Sigma, Cat#V2264) were resuspended in sterile water, whereas paclitaxel (Sigma, Cat#T7402), reversan (Sigma, Cat#SML0173), IWR-1 (Sigma, Cat#I0161) and LGK-974 (MedChemExpress, Cat#HY-17545) were dissolved in DMSO. Vehicle controls consisted of the corresponding solvent at matching final concentrations. Cell viability upon doxorubicin treatment in combination with reversan was assessed by crystal violet staining. Briefly, cells were fixed and stained with 0.5% (w/v) crystal violet prepared in 66% (v/v) methanol, followed by extensive washing with deionized water. Staining intensity was quantified by dissolving crystal violet in 1 ml of 10% (v/v) acetic acid for 15 min, after which 100 μl of the solution were transferred to a 96-well plate and optical density (OD) was measured at 570 nm using a GloMax Explorer microplate reader (GM3500; Promega). Viability was expressed as a percentage relative to vehicle-treated controls, set to 100%. For all other drug treatments, cell viability was assessed by real-time monitoring of cell confluence using an IncuCyte SX5 system (Sartorius), as described below.

### IncuCyte-based analysis of cell confluence and orange fluorescent object detection

Cells were seeded in 96-well plates at appropriate densities to ensure growth during the assay period. Twenty-four hours after plating, plates were transferred to the IncuCyte® SX5 Live-Cell Analysis System (Sartorius) to allow automated real-time imaging under standard culture conditions (37 °C, 5% CO₂). Phase-contrast and fluorescence images were acquired every 2 h using a 10× objective, with four non-overlapping fields captured per well over 48-72 h, as indicated for each experiment. Cell proliferation and viability were quantified by measuring confluence, defined as the percentage of the image area occupied by cells (highlighted with a colourful mask), using AI-based confluence segmentation implemented in IncuCyte 2023A software (Sartorius). For drug-response assays, cells were treated 24 h after seeding with increasing concentrations of doxorubicin, paclitaxel or vinorelbine. Growth kinetics were determined by plotting confluence over time, whereas dose-response curves were generated from endpoint confluence measurements normalized to vehicle-treated controls. IC₅₀ values were calculated by non-linear regression analysis using GraphPad Prism v9.5. For fluorescence-based analyses, orange fluorescent objects were automatically quantified using IncuCyte image analysis software with segmentation parameters optimized for the orange fluorescence channel. Background correction was performed using surface-fit background subtraction, and the object count unit (OCU) threshold was set to 0.1. Fluorescent signals were expressed as the number of orange objects per image, as indicated. All experiments were performed in at least three independent biological replicates.

### Generation of the constitutive active β-Catenin cells

A constitutively active form of β-Catenin was generated using the S33Y mutant construct (pcDNA3-β-Catenin S33Y)^45^, kindly provided by Eric Fearon (Addgene plasmid #19286). The empty pcDNA™3.1(+)/myc-His A vector (Invitrogen, Cat#2094) was used as control. Stable transfections were performed in TuBo-p140Cap cells using Lipofectamine 2000 (Thermo Fisher Scientific), according to the manufacturer’s instructions. TuBo-p140Cap cells were transfected with pcDNA3-β-Catenin S33Y, whereas TuBo-p140Cap and TuBo-Mock cells transfected with the empty vector served as controls. Following transfection, cells were selected in medium containing G418 (80 μg/ml; Thermo Fisher Scientific) for 14 days. Resistant colonies were isolated by manual picking and expanded in 96-well plates. Expression of constitutively active β-Catenin was confirmed by western blotting in our previous publication^29^. To reduce clonal variability, five independent β-Catenin-positive clones were pooled and used for all subsequent analyses.

### Transient silencing of ABCC1 in MDA-MB-231

Transient knockdown of ABCC1 was performed in MDA-MB-231 cells using ON-TARGET plus SMART pool small interfering RNA (siRNA; target gene: *ABCC1*, Entrez Gene ID: 4363; Dharmacon RNAi Technologies, GE Healthcare, Buckinghamshire, UK) or ON-TARGETplus non-targeting control siRNA (Dharmacon), following the manufacturer’s instructions. The ON-TARGET plus siRNA chemistry is designed to minimise off-target effects by chemical modification of both sense and antisense strands while preserving silencing efficacy. Cells were seeded in six-well plates and transfected at ∼70% confluence using Lipofectamine 2000 (Invitrogen, Thermo Fisher Scientific), according to the manufacturer’s protocol. After transfection, cells were maintained under standard culture conditions (37 °C, 5% CO₂) for 48 h before downstream analyses. Knockdown efficiency was assessed where indicated by quantitative PCR. Transfected cells were subsequently used in functional assays.

### RNA extraction, cDNA synthesis and qRT-PCR

Total RNA was isolated using TRIzol reagent (Invitrogen, Cat#15596018) according to the manufacturer’s protocol. RNA pellets were resuspended in nuclease-free water and stored at −80 °C. Prior to reverse transcription and quantitative PCR analysis, samples were thawed on ice. RNA concentration and purity were assessed using a NanoDrop™ One/OneC Microvolume UV-Vis Spectrophotometer (Thermo Fisher Scientific). Complementary DNA (cDNA) was synthesized from 1 μg of total RNA using random hexamer priming. qRT-PCR was performed using a 7900 Real-Time PCR System (Applied Biosystems), with reactions run in triplicate across at least three independent biological experiments. Relative gene expression was calculated using the 2^−ΔΔCT^ method, with normalization to the geometric mean of the housekeeping gene 18S rRNA to control for variability in input RNA and transcriptional efficiency. Primers used for qPCR are listed below:

Human *ABCC1*: CCGTGTACTCCAACGCTGACAT (Fw) – ATGCTGTGCGTGACCAAGATCC (Rv)

Human *ABCG2*: GTTCTCAGCAGCTCTTCGGCTT (Fw) – TCCTCCAGACACACCACGGATA (Rv)

Human *ABCB1*: GCTGTCAAGGAAGCCAATGCCT (Fw) – TGCAATGGCGATCCTCTGCTTC (Rv)

Human *ACTIN*: CACCATTGGCAATGAGCGGTTC (Fw) – AGGTCTTTGCGGATGTCCACGT (Rv)

Human *GAPDH*: GTGGAAGGGCTCATGACCA (Fw) – GGATGCAGGGATGATGTTCT (Rv)

Mouse *Abcc1*: CAGTGGTTCAGGGAAGGAGTCA (Fw) – CACTGTGGGAAGACGAGTTGCT (Rv)

Mouse *Abcg2*: CAGTTCTCAGCAGCTCTTCGAC (Fw) – TCCTCCAGAGATGCCACGGATA (Rv)

Mouse *Abcb1*: TCCTCACCAAGCGACTCCGATA (Fw) – ACTTGAGCAGCATCGTTGGCGA (Rv)

Mouse *Actin*: CATTGCTGACAGGATGCAGAAGG (Fw) – TGCTGGAAGGTGGACAGTGAGG (Rv)

### Droplet digital PCR

Droplet digital PCR (ddPCR) was performed for absolute quantification of target transcripts, particularly in samples with limited starting RNA input, using a droplet-based partitioning system (QX200™, Bio-Rad Laboratories, Hercules, CA, USA). Reactions were assembled in a final volume of 22 μl containing 11 μl of 2× ddPCR EvaGreen Supermix (Bio-Rad), forward and reverse primers at a final concentration of 100 nM each, 5 μl of template cDNA, and nuclease-free water. Droplets were generated using the QX200 droplet generator (Bio-Rad) by loading the reaction mix together with Droplet Generation Oil into DG8 cartridges (Bio-Rad), yielding approximately 15,000 droplets per sample. The resulting emulsions were transferred to a 96-well PCR plate, sealed, and subjected to thermal cycling under the following conditions: 95 °C for 5 min; 45 cycles of 95 °C for 30 s and 60 °C for 1 min; followed by a final enzyme deactivation step at 90 °C for 5 min. After amplification, droplets were stabilized for 30 min at 4 °C prior to fluorescence reading using the QX200 droplet reader (Bio-Rad). Data were analyzed using QX Manager Software (version 2.2 Standard Edition; Bio-Rad) according to the manufacturer’s instructions. The same primer sets used for qRT-PCR analyses were employed for ddPCR.

### Mammosphere Formation Assay

TuBo and 4T1 cells were seeded at a density of 6 × 10⁴ cells/ml in 10-cm ultra-low attachment dishes (Corning), or in Poly(2-hydroxyethyl methacrylate) (Poly-HEMA; Sigma-Aldrich) coated dishes (6-well or 24-well), using mammosphere serum-free DMEM/F12 medium (Invitrogen). The medium was supplemented with 20 ng/ml basic fibroblast growth factor (bFGF; Peprotech, Cat#100-18B), 20 ng/ml epidermal growth factor (EGF; Sigma-Aldrich, Cat#E9644), 5 μg/ml insulin (Sigma-Aldrich, Cat#I9278), 0.4% bovine serum albumin (BSA; Sigma), and 1% penicillin/streptomycin (Invitrogen). For MDA-MB-231 cells, the same plating density was used in 60-mm ultra-low attachment dishes in serum-free DMEM/F12 medium supplemented with 2% B27 supplement (Invitrogen, Cat#17504-044), 20 ng/ml EGF, 0.4% BSA, 4 μg/ml insulin, and 1% penicillin/streptomycin. After 5 days in culture, non-adherent spherical cell clusters (mammospheres) were analysed. For quantification, at least five representative fields per condition were acquired at day 5 using an IncuCyte SX5 live-cell imaging system (Sartorius) equipped with a 4× objective. Mammospheres were identified and quantified using size-based filtering parameters, with a minimum area threshold of 3,000 μm² and a maximum area threshold of 60,000 μm², and results were expressed as the number of mammospheres per well.

### Flow Cytometry (FC)

Cells were detached using 1% trypsin-EDTA (Gibco), collected, neutralized with complete DMEM, and washed twice with ice-cold phosphate-buffered saline (PBS). For analysis of intracellular doxorubicin accumulation, no additional staining was required due to the intrinsic fluorescence of doxorubicin (excitation, 488 nm; emission, 585/42 nm bandpass filter). Cells were immediately analysed by flow cytometry following resuspension in PBS. For apoptosis assays, cells were stained using the Annexin V Apoptosis Detection Kit (Invitrogen) according to the manufacturer’s instructions. Briefly, 1 × 10⁵ cells were resuspended in 100 μl of Annexin V binding buffer and incubated with 5 μl of eFluor™ 450-conjugated Annexin V for 15 min at room temperature in the dark. Samples were then washed and diluted in 300 μl of binding buffer and analysed immediately by flow cytometry. For cancer stem cell analysis in TuBo cells, single-cell suspensions were generated by trypsinization followed by neutralization with complete DMEM. Cells were centrifuged (5 min, 476 × g, 4 °C), resuspended in appropriate buffer, and counted. A total of 1 × 10⁶ cells per sample were incubated for 30 min at 4 °C in the dark with Sca-1-Alexa Fluor 647 antibody (BioLegend, Cat#122518; 1:200 dilution) in PBS. Flow cytometric acquisition was performed on a BD FACS Versе™ cytometer (BD Biosciences). A minimum of 30,000 events were recorded per sample. Gating strategies were established using unstained and/or single-stained controls, as appropriate. Doxorubicin-positive cells were identified based on fluorescence intensity in the Propidium Iodide (PI) channel, and data were analysed using BD FACSuite software (v1.3). All experiments were performed in at least three independent biological replicates.

### *In vivo* tumor growth

All in vivo experiments were performed using female mice only. Six- to eight-week-old female BALB/c mice were purchased from Charles River Laboratories (Calco, Italy) and maintained under specific pathogen-free conditions in accordance with institutional and European Community guidelines. For orthotopic tumour models, 1 × 10⁴ 4T1 cells or 1 × 10⁵ TuBo cells were resuspended in 50 μl of sterile PBS and injected into the left mammary fat pad of recipient BALB/c mice under aseptic conditions. Tumour growth was monitored every 2 days using a digital caliper by investigators blinded to experimental groups. Two perpendicular tumour diameters (length and width) were measured, and tumour volume was calculated using the ellipsoid formula: (4/3π(d/2)^2*D/2). Mice were euthanized when tumour volume reached approximately 600 mm³, in accordance with approved ethical limits; this threshold was not exceeded in any experiment. Throughout the study, animals were monitored regularly and did not exhibit weight loss exceeding 10% of baseline body weight. Doxorubicin hydrochloride (Doxo; Sigma-Aldrich, Cat#D1515) was freshly prepared in sterile water immediately prior to administration and injected according to the experimental design described in the relevant sections.

### *In vivo* treatments

Tumour-bearing mice were treated with doxorubicin hydrochloride (Doxo; Sigma-Aldrich, Cat#D1515) administered by intraperitoneal (i.p.) injection at doses of 3 mg/kg, 1.5 m/kg or 0.5 mg/kg, as indicated for each experiment. Treatments were administered every 3 days for a total of seven injections. Control animals received equivalent volumes of vehicle solution according to the same schedule. For WNT pathway inhibition studies, mice received intratumoral injections of IWR-1 (Sigma, Cat#I0161) at a dose of 5 mg/kg every 2 days for a total of seven administrations. Animals were monitored throughout the treatment period for tumour growth, body weight and signs of treatment-related toxicity.

### Tumor histological analysis

Tumors and lungs were fixed in 10% neutral buffered formalin and embedded in paraffin or fixed in 4% PFA and frozen in a cryo-embedding medium (OCT, Bio-Optica); 5 µm slides were cut and stained with Hematoxylin and Eosin (H&E, Bio-Optica) for histological examination. To optimize the detection of microscopic metastases and ensure systematic uniform and random sampling, lungs were cut transversally into 2.0-mm thick parallel slabs with a random position of the first cut in the first 2 mm of the lung, resulting in 5-8 slabs for lungs. The slabs were then embedded cut surface down and sections were stained with H&E. Slides were independently evaluated by two pathologists to quantify the number of metastatic lesions.

For immunohistochemistry on frozen samples, sections were air-dried, fixed in ice-cold acetone for 10 min, and incubated with rat monoclonal anti-CD31 (550274, BD Pharmingen) mixed with rat monoclonal anti-CD105 (550546, BD Pharmingen) antibodies, followed by incubation with the secondary antibody (Jackson ImmunoResearch Laboratories). Immunoreactive antigens were detected using alkaline phosphatase-conjugated streptavidin (Thermo Scientific) and Vulcan fast red Chromogen (Biocare Medical). After chromogen incubation, slides were counterstained in Hematoxylin (Bio-Optica), and images were acquired using a Leica DMRD optical microscope.

Immunohistochemical staining on formalin fixed paraffin-embedded (FFPE) tissues was performed using the following primary antibodies: mouse monoclonal anti-γH2AX (Millipore) and rabbit anti-mouse cleaved Caspase-3 (R&D System). Antigen retrieval was performed by microwaving slides for 10 min in Tris-EDTA buffer (pH 9.0, Sigma Aldrich). Envision anti-rabbit or anti-mouse IgG (Dako) secondary antibodies were applied for 30 min, followed by signal development using 3-3’ diaminobenzidine (DAB) as the chromogen. After counterstaining with Hematoxylin (Bio-Optica), images were scanned with Nanozoomer scanner from Hamamatsu. The number of γ-H2AX positive cells and the percentage of Caspase3 positive cells were calculated with Qu-Path 0.3.2 software using positive cell detection tool.

For immunofluorescence, 4-6 µm cryostat sections were air-dried, fixed in ice-cold acetone for 10 min and incubated with the following primary antibodies: rat monoclonal anti-CD31 (550274, BD Pharmingen) mixed with rat monoclonal anti-CD105 (550546, BD Pharmingen), rabbit polyclonal anti-NG2 (ab5320, Millipore) and mouse monoclonal anti-αSMA (M0851, Dako), followed by secondary antibodies conjugated with Alexa 546 and Alexa 488 (Invitrogen). Nuclei were stained with DRAQ5 (Life Technologies). Images were acquired using Zeiss LSM 510 META confocal microscope. Quantification of vascular area and pericyte coverage was performed on the digital images of 11-14 tumors per group (4 X 200 microscopic fields per tumor) by 2 pathologists, independently and in a blind fashion, using Adobe Photoshop.

### Dataset analysis

Analysis of transcriptomic data from BC patient cohorts was performed using the METABRIC and TCGA datasets. Single-sample gene set enrichment analysis (ssGSEA) was conducted in R using the GSVA package to assess pathway enrichment at the individual patient level. Correlations between pathway enrichment scores and ABCC1 expression were calculated and visualized using the ggpubr package in R. Additional analyses were performed using publicly available BC datasets and web-based platforms, including ROCplot.org^30^ for therapy-response analyses, integrated clinical and molecular datasets from METABRIC, TCGA and IMPACT^31^ for survival analyses, and GEPIA3^38^ for transcriptomic profiling and gene expression analyses. Analyses were carried out according to the corresponding platform guidelines and default settings, unless otherwise specified. Publicly available normalized proteomic datasets were further interrogated to assess the association between p140Cap and ABCC1 protein expression. For primary TNBC patients, normalized proteomic data from the Dan L Duncan Comprehensive Cancer Center (DLDCCC) cohort were obtained from the study by Anurag et al^54^. Pearson correlation analyses were performed to evaluate the relationship between p140Cap and ABCC1 protein across datasets. In addition, pre-processed single-cell RNA-seq data from two triple-negative breast cancer patients were retrieved from GEO (GSE263995; 22,487 cells before and after chemotherapy)^55^. Basal-like epithelial cells were identified by excluding stromal/immune/endothelial cells based on canonical markers (including PTPRC, PECAM1, PDGFRA, PDGFRB and ACTA2) and by selecting cells enriched for basal-like markers (including KRT5, KRT6A/B/C, KRT14, KRT15, KRT16, KRT17, TP63, SOX10, FOXC1, CDH3, EGFR, ITGA6, ITGB4 and PROM1), yielding 1,013 basal-like epithelial cells. These cells were then used to assess intratumoral transcriptional heterogeneity and SRCIN1/ABCC1 expression patterns. Based on normalized expression values, cells were classified as SRCIN1+/ABCC1−, ABCC1+/SRCIN1−, co-expressive or non-expressive. Spearman correlation analysis was performed on cells expressing at least one of the two genes (n = 121), revealing a significant inverse association between SRCIN1 and ABCC1 expression in basal-like epithelial cells (Rho = −0.56, p = 1.5 × 10-11).

### PDX validation analysis

Pt A, Pt B and Pt C are three independent patient-derived TNBC cell populations, previously stratified for p140Cap expression by immunohistochemical (IHC) analysis, cultured in 1:1 mixture of DMEM and Ham’s F12 medium, supplemented with 2 mM L-Glutamine, 5 μg/ml insulin, 0.5 μg/ml hydrocortisone, 2% B27, 20 ng/ ml EGF and FGF, and 4 μg/ml heparin.

The lentiviral construct harboring the p140Cap protein was engineered by VectorBuilder, subcloning the full-length human p140Cap cDNA [NM_025248.3] of pLVX puro lentiviral vector. These vectors were used for stable overexpress p140Cap. The pLVX empty vector served as a negative control. PDX cell lines were stably transduced with the lentiviral vectors in the presence of 4 μg/mL of polybrene (Sigma). After 48 h, the infected cells were placed in fresh medium with 2 μg/mL of puromycin (Vinci Bio Chem) and selected for 72 h.

#### PDX immunohistochemistry

Three-μm thick sections were prepared from FFPE tissue blocks, dried at 37 °C O/N and then processed with Bond-RX fully automated stainer system (Leica Biosystems) according to the following protocol. First, tissues were deparaffinized and pretreated with the Epitope Retrieval Solution 1 (pH 9) at 100°C for 20 min. After the washing steps, peroxidase blocking was performed for 10 min using the Bond Polymer Refine Detection Kit (#DC9800; Leica Biosystems). Tissues were incubated for 30 min with the anti-p140Cap Ab diluted 1:1000 in Bond Primary Ab Diluent (#AR9352). Subsequently, tissues were incubated with post primary and polymer for 16 min, developed with DAB-chromogen for 10 min, and counterstained with hematoxylin for 5 min. Slides were digitally scanned with the Aperio ScanScope.

#### PDX quantitative real-time PCR

RT-qPCR was performed using the Ssofast EvaGreen SYBR Supermix ® (BIO-RAD). The ΔCt method was used to calculate the mRNA levels of each target gene normalized against different housekeeping genes. The 2^−ΔΔCt^ method was used to compare the mRNA levels of each target gene and the relative amplification value was plotted in the graph. The primer sequences were used for SYBR Green methodology.

#### PDX immunofluorescence studies

PDX cells were seeded on poly-D-lysine coated glass slides, left to adhere for 10 min and fixed with 4% parformaldehyde (20 min at RT). Fixed cells were permeabilized in PBS 0.1% Triton X-100 for 5 min at RT. To prevent non-specific binding of the Ab, cells were incubated with PBS in the presence of 3% BSA for 30 min. Immunostaining was carried out with anti-p140Cap antibody (1.6 mg/ml) for 2 hours at RT in blocking solution, followed by incubation with donkey-a-mouse (AlexaFluor Plus 488, Thermofisher, 1:400) secondary antibody for 1 hour at RT. After three washes with PBS, nuclei were DAPI stained for 30 min at RT, and after three washes in PBS, the coverslips were mounted in Mowiol-Dabco.

Images were acquired using a Leica SP8 Confocal microscope equipped with a 63X/1.4NA oil-immersion objective and data were collected with LasX (Version 3.5.5 Leica). Brightness and contrast optimizations were applied uniformly across the entire image for visualization purpose only. Percentage of p140Cap-overexpressing cells was quantified in Fiji (v1.52, National Institutes of Health) by setting a threshold on control samples such that just the 0.05% of pixels were positive, thereby excluding the endogenous signal. The same cutoff was applied on images of p140Cap-overexpressing cells. Based on this criterion, cells were manually classified as positive or negative for protein overexpression. Quantification was performed on n > 80 cells per condition.

#### PDX doxorubicin accumulation analysis

Cellular accumulation of doxorubicin was assessed exploiting the intrinsic fluorescence properties of the compound. Briefly, PDX cells were seeded in 24-well plates at a density of 5 × 10⁴ cells/well and cultured under standard conditions. After 72 h, cells were incubated with 1 μM doxorubicin in complete culture medium for 1 h at 37°C. Following treatment, cells were transferred onto poly-D-lysine-coated glass coverslips, allowed to adhere for 10 min, and fixed with 4% paraformaldehyde for 20 min at room temperature (RT). Cells were then permeabilized with 0.1% Triton X-100 in PBS for 5 min at RT. Nuclei were stained with DAPI for 30 min at RT. After three washes with PBS, coverslips were mounted using Mowiol-DABCO mounting medium. Fixed cells were permeabilized in PBS 0.1% Triton X-100 for 5 min at RT. Nuclei were DAPI stained for 30 min at RT, and after three washes in PBS, the coverslips were mounted in Mowiol-Dabco. Doxorubicin fluorescence was visualized using a Leica DM6 B MultiFluo equipped 20X / 0.75 NA dry objective. Images were acquired with LasX software using identical exposure settings across all experimental conditions to allow quantitative comparison among samples (doxorubicin excitation/emission: ∼540/592 nm). Quantitative analysis of intracellular doxorubicin accumulation was performed using Fiji software. Cell nuclei were segmented based on the DAPI channel using StarDist in Fiji, and the mean doxorubicin fluorescence intensity within each nuclear region was measured. For each sample, fluorescence values were normalized by dividing them by the median fluorescence intensity of the corresponding control empty vector (E.V.), and data are presented as relative mean doxorubicin fluorescence.

### Statistical analysis and reproducibility

Statistical analyses were performed using GraphPad Prism v9.5.0 (GraphPad Software). Unless otherwise indicated, data are presented as mean ± s.e.m. (SEM), and individual data points are shown when appropriate to illustrate data distribution. Data distribution was assumed to be normal, although this was not formally tested. Statistical significance between two groups was determined using unpaired two-tailed Student’s *t*-tests unless otherwise specified in the figure legends or relevant Methods sections. Kaplan-Meier survival analyses were used to estimate overall survival, and statistical significance was evaluated. Exact *P* values are reported in the figures or corresponding legends, and *P* < 0.05 was considered statistically significant. IC_50_ values were calculated by non-linear regression analysis using the default dose-response curve fitting model implemented in GraphPad Prism. Correlation analyses were performed using Pearson’s correlation coefficient unless otherwise indicated. Experiments were independently repeated as specified in the figure legends to ensure reproducibility. Sample sizes were not predetermined by formal statistical methods but were consistent with those commonly used in the field and in our previous related studies. No data were excluded from the analyses. Cell-based experiments were not randomized, and investigators were generally not blinded to group allocation or outcome assessment, unless otherwise stated. In vivo tumour measurements were performed in a blinded manner.

## Code availability

This study did not involve the generation or use of custom code.

## Acknowledgements

We are grateful to Francesca Dionisio for her expert guidance in the introduction and optimization of the ddPCR methodology. We thank Deborah Traversi for generously providing access to the QX200 system (Bio-Rad) at the Department of Public Health and Paediatric Sciences, University of Turin. We also acknowledge Enrico Patrucco for his valuable support and training in the use of the Incucyte SX5 (Sartorius) at the Molecular Biotechnology Center (MBC), University of Turin. The authors received funding from Mur, PRIN 2022 Project number 2022WYAEWE PI Defilippi Paola; from AIRC under IG 2022 - ID. 27353 project - P.I. Defilippi Paola. This work was also supported by: Fondazione CRT 2020.1798, RILO University of Torino (IG 11904, IG 15538), Ministero della Salute (RF-2021-12371961) to Paola Defilippi, PNRR M4C2-Investimento 1.4-529 CN00000041 “Finanziato dall’Unione Europea-NextGenerationEU” to Paola Defilippi and Daniela Taverna. All schematic illustrations included in this manuscript were created with BioRender.com.

## Author information

### Contributions

A.S., M.P., P.D. and V.S. conceptualized and designed the study. A.S., M.P., A.SAR. and V.S. performed the in vitro and in *vivo* experiments and acquired the data. A.L., M.I. processed and performed IHC analysis of murine tumor samples. S.P., D.TOS., M.G.F., and L.B. analyzed the human BC samples. A.S., V.S., M.J., U.A., Y.V., S.L. and F.A.T. performed the bioinformatics analysis. A.S, M.P. and V.S. analyzed and interpreted all the data. A.S. and V.S drafted the paper. All authors reviewed and revised the paper. V.S., A.S., E.T., D.TAV., F.O., B.B. and P.D. provided financial support, continuous data review through regular meetings and the active pursuit of collaborations with external colleagues. V.S. and P.D. supervised the study.

## Extended data

**Extended Data Fig. 1:**
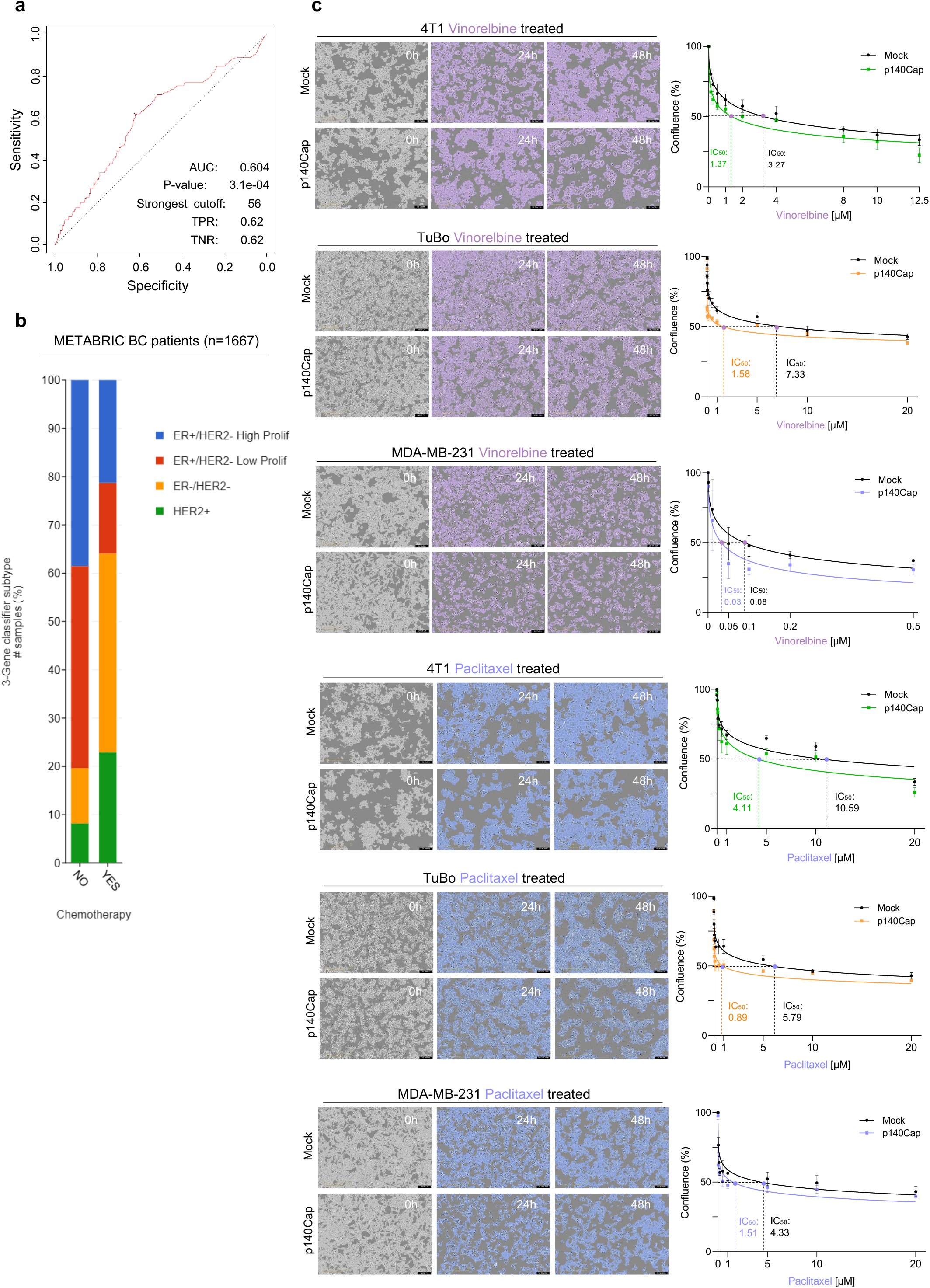
p140Cap increases sensitivity to vinorelbine and paclitaxel. **a**, ROC curve between SRCIN1 expression levels and chemotherapy response in chemotherapy-treated BC patients^33^. The area under the curve (AUC) was 0.604 (*P* = 3.1 × 10⁻⁴). The strongest cutoff for SRCIN1 expression was 56, yielding a true positive rate (TPR) of 0.62 and a true negative rate (TNR) of 0.62. **b**, Bar plot showing the distribution of luminal, HER2⁺, and TNBC molecular subtypes within the METABRIC cohort (*n* = 1,667 BC samples) chemotherapy treated (YES) or non-treated (NO). Bars indicate the percentage of cases assigned to each subtype based on 3-Gene classifier. **c**, Representative live-cell images (left) and dose-response curves (right) of Mock and p140Cap 4T1 (images displayed, Vinorelbine: 1 µM, Paclitaxel: 10 µM), TuBo (images displayed, Vinorelbine: 0.1 µM, Paclitaxel: 0.5 µM) and MDA-MB-231 (images displayed, Vinorelbine: 0.5 µM, Paclitaxel: 1 µM) cells treated with Vinorelbine and Paclitaxel. Cells were exposed to increasing concentrations of each drug (0-20 µM) and monitored for 0-48 h using the IncuCyte SX5. Quantitative analysis was performed with the Cell-by-Cell analysis module; analysis masks are shown in grey (pre-treatment), purple (post-vinorelbine) and light blue (post-paclitaxel). IC_50_ curves were generated from dose-resolved measurements of cell confluence. Data represent biological triplicates.

**Extended Data Fig. 2:**
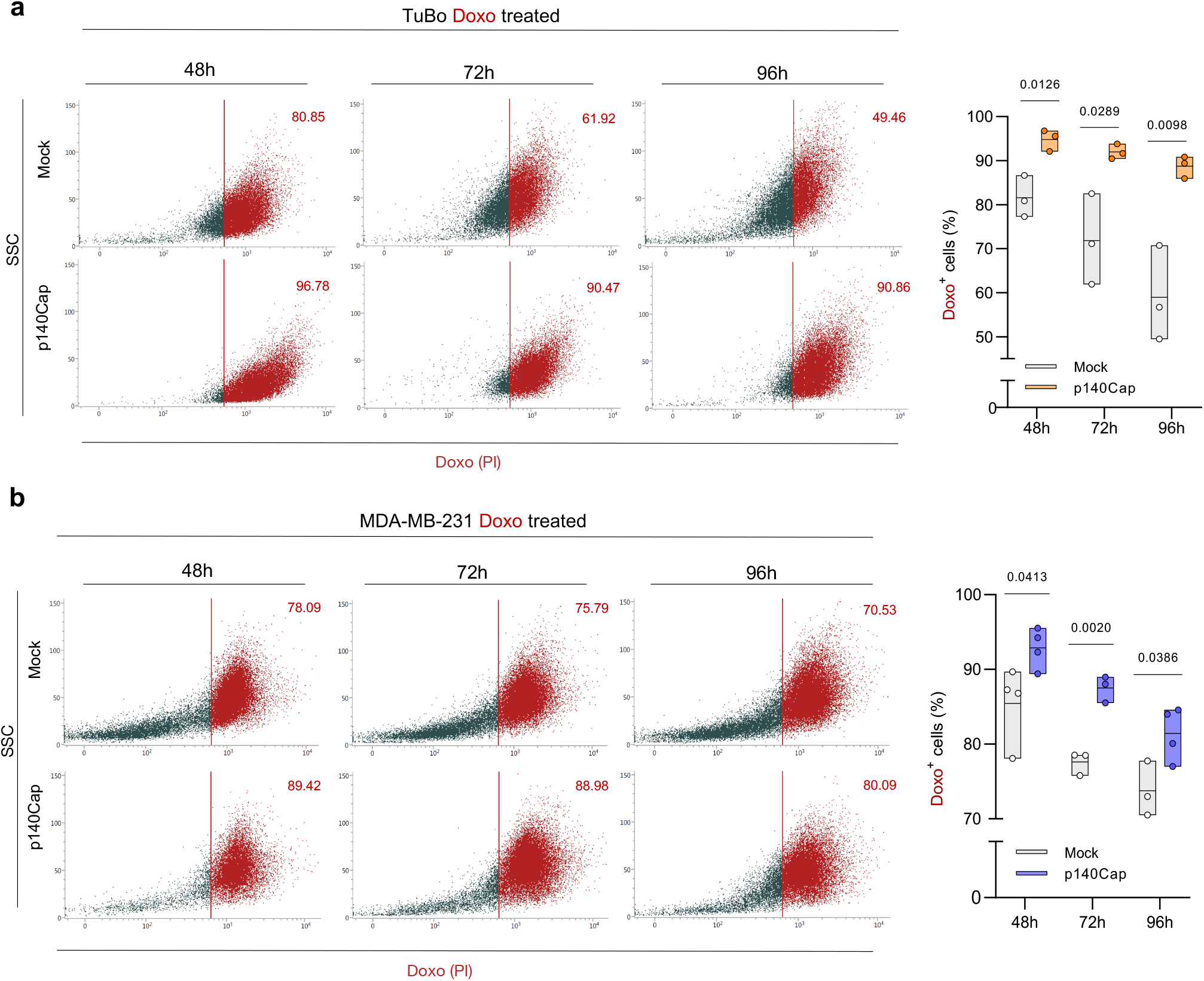
Sustained intracellular retention of doxorubicin in p140Cap cells. **a, b,** Representative flow cytometry plots and floating bar (min-to-max) quantification of Doxo-positive cells in Mock and p140Cap TuBo and MDA-MB-231 cells. Cells were treated with 0.125 µM and 0.5 µM Doxo, respectively, and analyzed 48/72/96 h post-treatment. Gating was established based on the corresponding untreated control. Data derived from three independent experiments.

**Extended Data Fig. 3:**
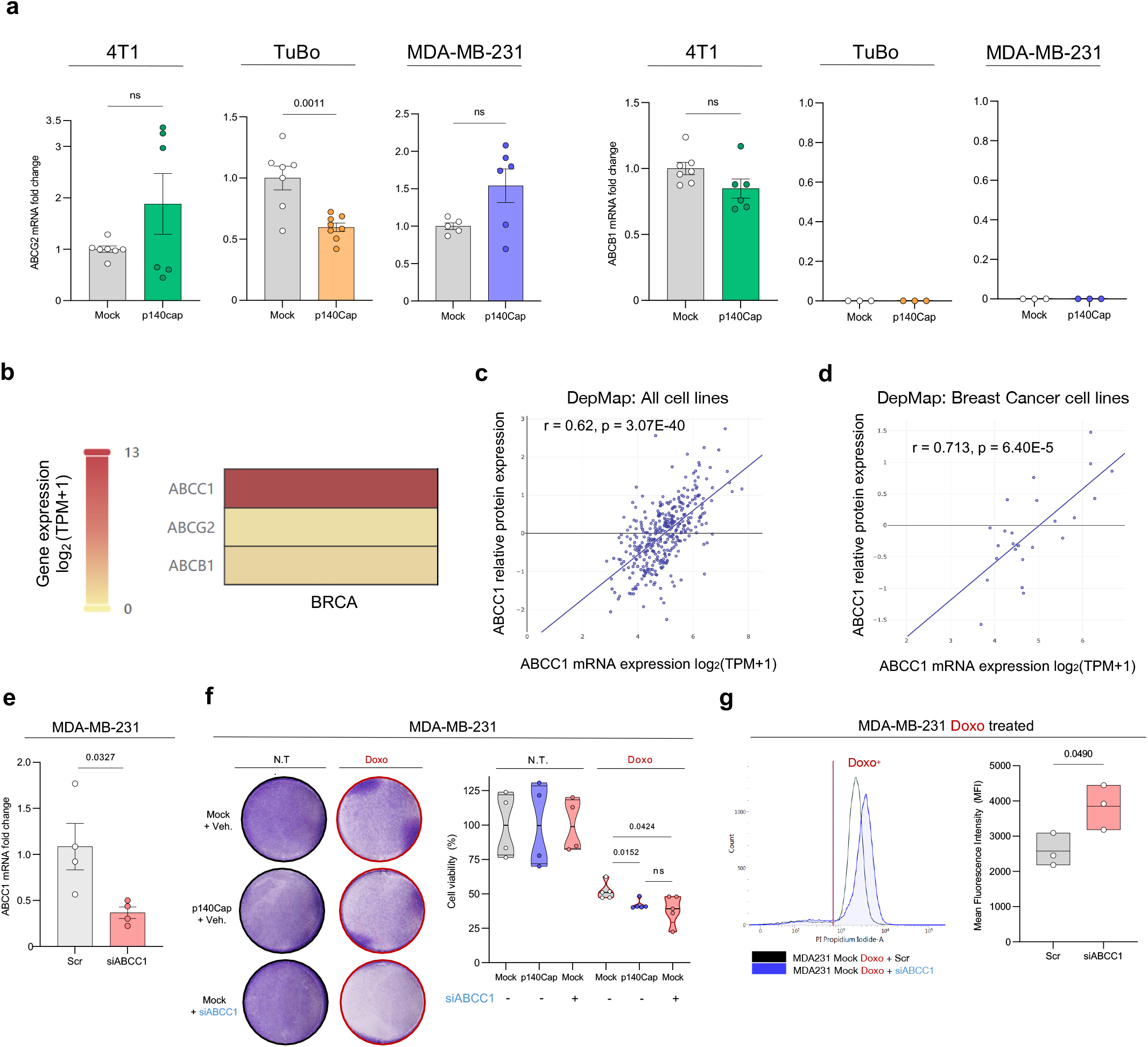
Functional validation of ABCC1 in doxorubicin sensitivity in BC. **a**, qRT-PCR analysis of ABCG2 (left) and ABCB1 (right) in Mock and p140Cap 4T1, TuBo and MDA-MB-231 cells. Relative mRNA expression levels were normalized to 18S rRNA and expressed as fold change relative to ABCC1 Mock controls. Data are presented as mean ± SEM from biological replicates (dots), with statistical analysis performed using unpaired t-tests. **b**, Heatmap showing normalized expression levels of *ABCC1* across breast cancer (BRCA) samples from TCGA, generated using GEPIA3^41^. Expression levels are scaled and color-coded from low (yellow) to high (red) based on log_2_ (TPM + 1). **c**, **d**, Scatter plots show the correlation between ABCC1 mRNA log_2_ (TPM + 1) and protein (sp|P33527|) expression levels across All cell lines (b) and Breast Cancer cell lines (c) from the DepMap dataset. Correlation coefficients were calculated using Pearson’s correlation analysis, with two-sided *P* values indicated in the plot. **e**, Validation of ABCC1 silencing in MDA-MB-231 Mock cells by qRT-PCR. ABCC1 mRNA levels were normalized to 18S rRNA and expressed relative to control cells. Data are shown as mean ± SEM from 4 biological replicates. **f**, Representative Crystal Violet staining (left) and quantification (right) of cell viability in MDA-MB-231 cells treated with Doxo 1 µM in the presence or absence of siABCC1. Crystal Violet staining was quantified as a measure of relative cell viability to N.T. counterparts. Data represent *n* = 5 biological replicates. **g**, Representative flow cytometry histograms (left) and mean fluorescence intensity (MFI) quantification (right) of MDA-MB-231 Mock cells treated with 0.5 µM Doxo in the presence or absence of siABCC1. Data represent biological triplicates, with statistical analysis performed as described in the Methods.

**Extended Data Fig. 4:**
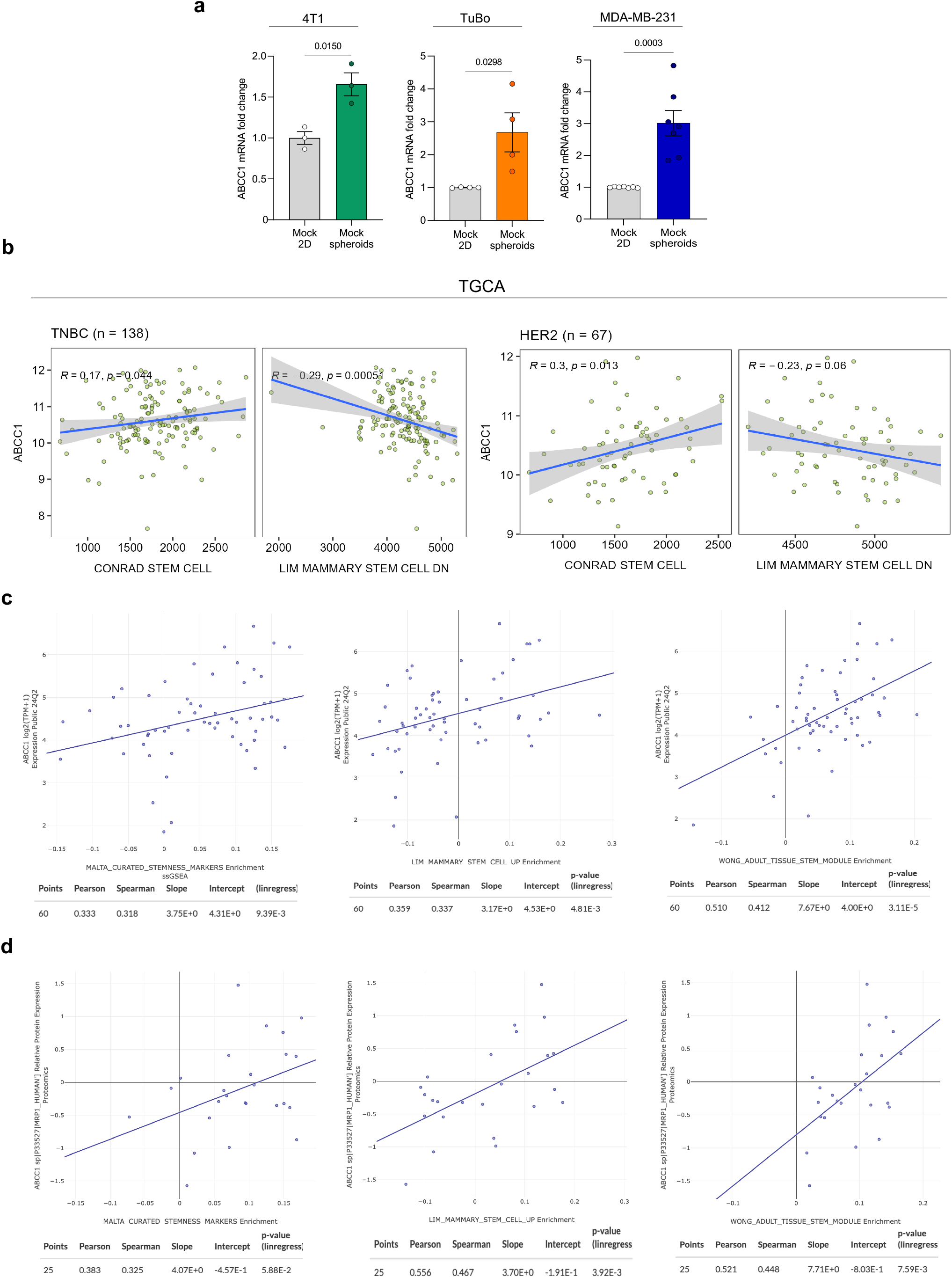
ABCC1 expression is enriched in mammospheres and correlates with stemness signatures in clinical datasets. **a**, qRT-PCR analysis of ABCC1 expression in Mock 2D monolayer versus mammosphere cultures of 4T1 (*n* = 3), TuBo (*n* = 4), and MDA-MB-231 (*n* = 7) cells. Expression levels were normalized to 18S rRNA and are shown relative to Mock 2D cultures. Data represent biological replicates and are presented as mean ± SEM. **b-d** Scatter plots showing the correlation between ABCC1 and the indicated gene sets in TNBC patients (n = 138 patients) and HER2 patients (n = 67 patients), data taken from the TCGA dataset (b), and in breast cancer cell lines from the DepMap resource (c,d). Pearson correlation coefficients (*r*) and two-sided *P* values are indicated.

**Extended Data Fig. 5.**
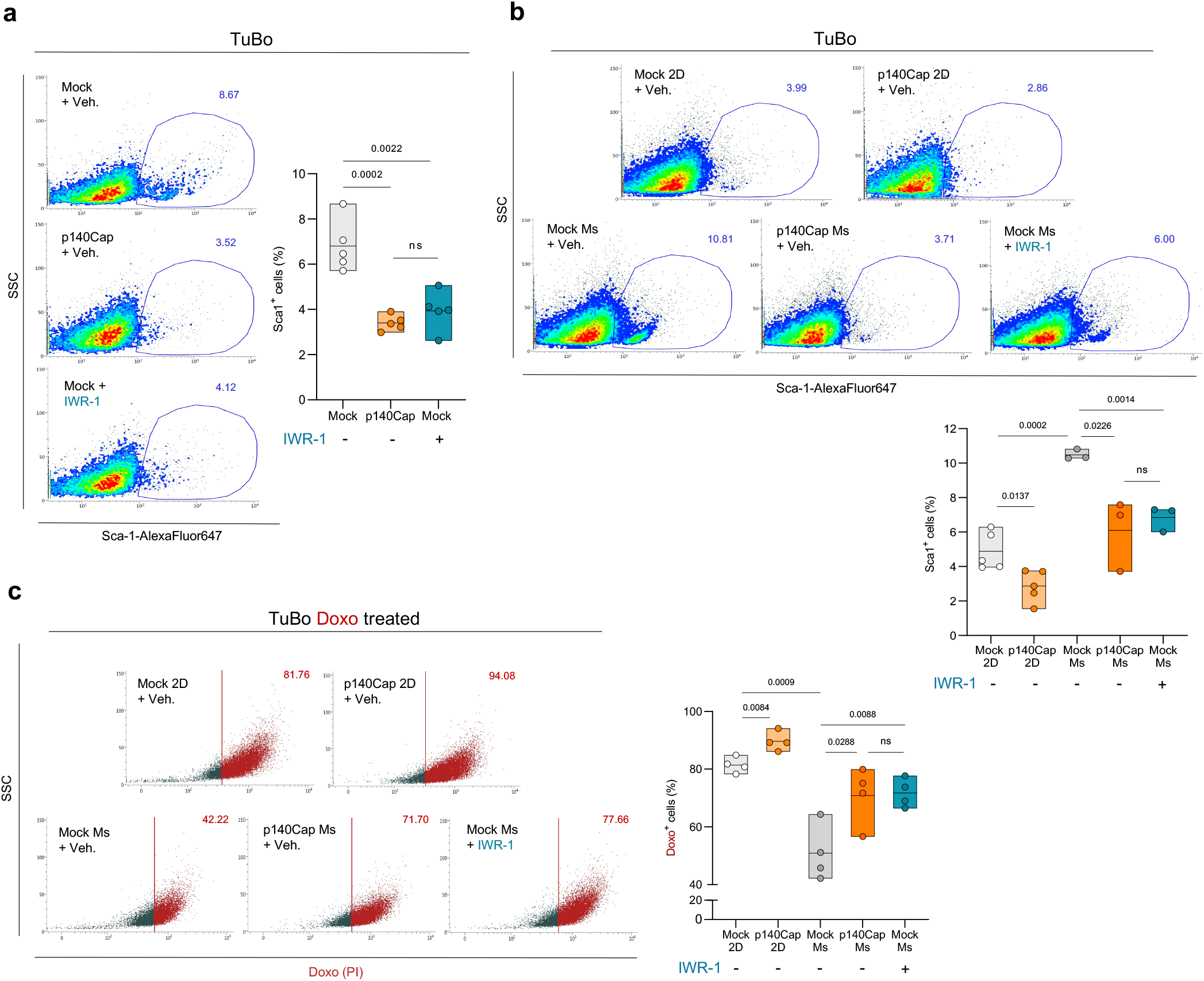
IWR-1 reduces stem-like populations. **a**, qRT-PCR analysis of ABCC1 expression in Mock 2D monolayer versus mammosphere cultures of 4T1 (*n* = 3), TuBo (*n* = 4), and MDA-MB-231 (*n* = 7) cells. Expression levels were normalized to 18S rRNA and are shown relative to Mock 2D cultures. Data represent biological replicates and are presented as mean ± SEM. **b-d** Scatter plots showing the correlation between ABCC1 and the indicated gene sets in TNBC patients (*n* = 138 patients) and HER2 patients (*n* = 67 patients), data taken from the TCGA dataset (b), and in breast cancer cell lines from the DepMap resource (c,d). Pearson correlation coefficients (*r*) and two-sided *P* values are indicated.

**Extended Data Fig. 6:**
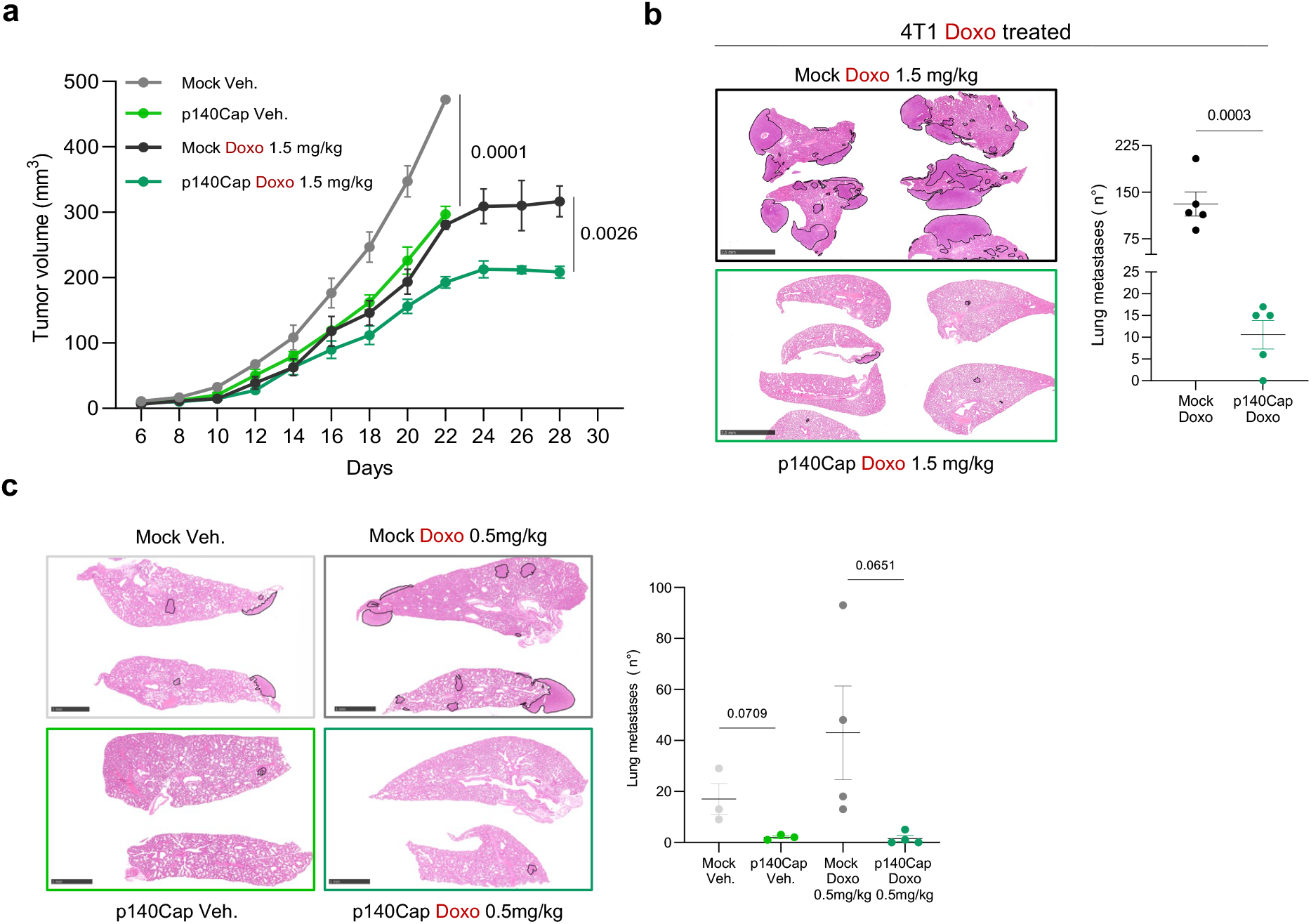
p140Cap reduces 4T1 tumor growth and metastatic lesions upon doxorubicin treatment. **a,** Tumor growth curves of orthotopic 4T1 Mock and p140Cap tumors treated with vehicle (Veh.) or Doxo 1.5 mg/kg, as indicated. Tumor volumes were measured at the indicated time points after injection. Data are mean ± SEM. Exact *P* values are shown in the panel. **b**, Representative images of hematoxylin and eosin (H&E) staining of lung sections from 4T1 tumor-bearing mice treated with Doxo. Metastatic lesions are outlined in black. Scale bars, as shown. Right, quantification of the number of lung metastases per mouse. Dots represent individual mice and bars indicate mean ± SEM. Statistical analysis was performed using a two-tailed unpaired *t*-test. **c**, Representative images of H&E-stained lungs and quantification of lung metastases in 4T1-bearing mice from the indicated groups. Metastatic lesions are outlined in black. Scale bars, as shown.

## Supplementary information

**Supplementary video 1**: Time-lapse of intracellular doxorubicin accumulation (red dots) and cell confluence of TuBo Mock cells. Cells were exposed to 1 μM Doxo for 6 h followed by drug washout, and monitored using an IncuCyte live-cell imaging system. Doxo accumulation was quantified every 2 h for 48 h following treatment initiation, by automated orange object count implemented in Incucyte software. TuBo Mock cells present a more efficient intracellular doxorubicin clearance which in turn reduces the drug’s cytotoxic efficacy. Data are representative of three biologically independent experiments.

**Supplementary video 2**: Time-lapse of intracellular doxorubicin accumulation (red dots) and cell confluence of TuBo p140Cap cells. Cells were treated and analysed as Mock cells. TuBo p140Cap cells show higher doxorubicin retention, which enhances chemosensitivity. Data are representative of three biologically independent experiments.

